# Quantitative mapping of fluorescently tagged cellular proteins using FCS-calibrated four dimensional imaging

**DOI:** 10.1101/188862

**Authors:** Antonio Z. Politi, Yin Cai, Nike Walther, M. Julius Hossain, Birgit Koch, Malte Wachsmuth, Jan Ellenberg

## Abstract

**EDITORIAL SUMMARY:** This protocol describes how to estimate and spatially resolve the concentration and copy number of fluorescently tagged proteins in live cells using fluorescence imaging and fluorescence correlation spectroscopy (FCS).

**TWEET:** Determining protein concentrations and copy numbers in live cells using fluorescence correlation spectroscopy (FCS)-calibrated imaging.

**COVER TEASER Map protein concentrations with FCS-calibrated imaging:** **Up to four primary research articles where the protocol has been used and/or developed:**

1. Walther, N., Hossain, M. J., Politi, A. Z., Koch, B., Kueblbeck, M., Oedegaard-Fougner, O., Lampe, M. and J. Ellenberg (2018). A quantitative map of human Condensins provides new insights into mitotic chromosome architecture. *bioRxiv*, 237834. https://doi.org/10.1101/2378342.
2. Cai, Y., Hossain, M. J., Heriche, J.-K., Politi, A. Z., Walther, N., Koch, B., Wachsmuth, M., Nijmeijer, B., Kueblbeck, M., Martinic, M., Ladurner, R., Peters, J.M. and J. Ellenberg (2017). An experimental and computational framework to build a dynamic protein atlas of human cell division. *bioRxiv*, 227751 https://doi.org/10.1101/227751
3. Germier, T., Kocanova, S., Walther, N., Bancaud, A., Shaban, H.A., Sellou, H., Politi, A.Z., Ellenberg, J., Gallardo, F. and K. Bystricky (2017). Real-Time Imaging of a Single Gene Reveals Transcription-Initiated Local Confinement. *Biophysical Journal*, 113(7), 1383-1394, https://doi.org/10.1016/j.bpj.2017.08.014.
4. Cuylen, S., Blaukopf, C., Politi, A. Z., Muller-Reichert, T., Neumann, B., Poser, I., Ellenberg, J., Hyman, A.A., and D.W. Gerlich (2016). Ki-67 acts as a biological surfactant to disperse mitotic chromosomes. *Nature*, 535(7611), 308–312. http://doi.org/10.1038/nature18610.

**Abstract:** The ability to tag a protein at its endogenous locus with a fluorescent protein (FP) enables the quantitative understanding of protein dynamics at the physiological level. Genome editing technology has now made this powerful approach routinely applicable to mammalian cells and many other model systems, opening up the possibility to systematically and quantitatively map the cellular proteome in four dimensions. 3D time-lapse confocal microscopy (4D imaging) is an essential tool to investigate spatial and temporal protein dynamics, however it lacks the required quantitative power to make absolute and comparable measurements required for systems analysis. Fluorescence correlation spectroscopy (FCS) on the other hand provides quantitative proteomic and biophysical parameters such as protein concentration, hydrodynamic radius and oligomerization but lacks the ability for high-throughput application in 4D spatial and temporal imaging. Here, we present an automated experimental and computational workflow that integrates both methods and delivers quantitative 4D imaging data in high-throughput. These data is processed to yield a calibration curve relating the fluorescence intensities of image voxels to absolute protein abundance. The calibration curve allows the conversion of the arbitrary fluorescence intensities to protein amounts for all voxels of 4D imaging stacks. With our workflow the users can acquire and analyze hundreds of FCS-calibrated image series to map their proteins of interest in four dimensions. Compared to other protocols, the current protocol does not require additional calibration standards and provides an automated acquisition pipeline for FCS and imaging data. The protocol can be completed in 1 day.

## INTRODUCTION

Tagging of proteins with fluorescent markers is essential to study their localization, function, dynamics and interactions in living cells. With the advantage of genome editing technologies such as CRISPR/Cas9 ^1–3^, it is now possible to engineer almost any higher eukaryotic cell type to homozygously express a protein of interest (POI) fused to a FP at its physiological level (see our accompanying Nature Protocol ^4^). Genome editing in combination with absolute quantitative fluorescence microscopy extends proteomics methods, which are typically restricted to large cell populations at specific time points, to single cell dynamic proteomics of living cells.

Modern confocal microscope detectors show a linear dependency of fluorophore concentrations and fluorescence intensities within several orders of magnitudes. To perform absolute quantitative imaging and convert relative fluorescence intensities to absolute physical quantities, such as the concentration of the fluorophore bearing protein, one can determine the calibration parameters of this linear dependency. To estimate the fluorophore’s concentration, temporal and spatial fluctuations of diffusing species can be used ^5–7^ In this protocol we describe how to use single point confocal FCS, a biophysical single molecule technique that allows to measure concentrations and diffusion coefficients of fluorescently labeled molecules ^7–10^. This method is well suited for low to medium abundant proteins (concentration range from few pM up to μM), a situation often observed at endogenous expression levels. Furthermore, most of the commercially available confocal microscopes can be equipped with the necessary detectors and hardware to perform FCS, making FCS-calibrated quantitative imaging easily available. In this protocol, FCS is combined with 3D confocal time-lapse imaging to estimate the calibration parameters and convert fluorescence intensities to concentrations in living cells. We previously used this approach to estimate protein numbers on bulk chromatin ^11,12^, on chromosome boundaries ^13^, large multi-protein complexes ^14^, and throughout mitosis for several genome edited cell lines ^12,15^. In this protocol, we offer custom software packages and a step-by-step guide based on a Zeiss LSM confocal microscope with FCS capability as a widely used commercial system. The data acquisition software packages (available in **Supplementary Software 1-3**) simplify the repeated cycles of imaging and FCS measurements required for FCS-calibrated imaging. In combination with the provided online image analysis tools (**Box 1**), data acquisition can be completely automated ^16–18^. This allows, without human supervision and in high-throughput, to automatically select cells with the optimal protein expression level and morphology and acquire images and FCS measurements. The data analysis software package (**Supplementary Software 4**) can be used to automatically extract the fluorescence intensities at FCS measurement points, compute protein concentrations and the FCS calibration parameters, and convert fluorescence image intensities to protein concentrations and absolute protein numbers.

## FLUORESCENCE CORRELATION SPECTROSCOPY

For a detailed description on FCS and dual color fluorescence cross-correlation spectroscopy (FCCS) we refer readers to reviews and protocols (e.g. ^6–10^). Here we provide only a short and practically oriented introduction to the topic (Fig. 1). FCS measures the fluorescence intensity (FI) fluctuations of fluorescent molecules in a sub-femtoliter observation volume of a focused laser beam. The intensity changes caused by single molecules moving in and out of the observation volume over time are recorded over a time-frame of 15-30 sec (Fig. 1a-b). The FI fluctuations in the observation volume depend on *(i)* the concentration of fluorophores, *(ii)* their mobility, *(iii)* their photophysics, and *(iv),* to a lesser extent, the properties of the detector. To analyze the fluctuations, the autocorrelation function (ACF) *G(τ)* is computed from the intensity time trace *I/(t) = ⟨I⟩ + δI(t)*:

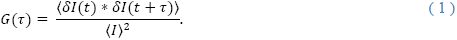

The brackets ⟨·⟩ indicate time-averaged quantities and *δI(t)* the mean of the time-averaged intensity *⟨I⟩*. The amplitude of *G(τ)* is inversely proportional to the number of particles in the observation volume *N* (Fig. 1c). The decay time *τ_D_* of *G(τ)* gives the characteristic time a fluorophore remains in the observation volume and, consequently, is a direct measure of the fluorophore diffusion. These parameters can be extracted by fitting the ACF to a physical diffusion model (see **Supplementary Notes 1** and **2**). To compute the concentration of the fluorophore at the FCS measurement point, the number of molecules *N* is divided by an effective confocal volume *V_eff_* (**Box 2**), typically below 1 fl, measured for the used microscopy setup.

**Figure 1:**
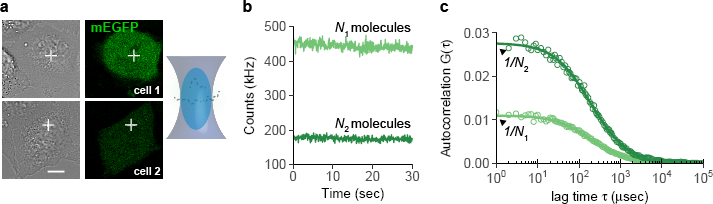
Principle of FCS for point confocal microscopes. (a) The excitation laser beam is positioned to a specific location of a cell (white cross). Fluorophores entering the confocal volume are excited and the number of photons emitted is recorded. Shown are two cells expressing different levels of the fluorescent protein mEGFP. Scale bar 10 μm.
(b) The fluctuations of the fluorophores diffusing in and out of the confocal volume cause fluctuations in the number of photons.
(c) Computing the self-similarity of the fluctuations in (b) yields ACFs. The amplitude of the ACF is directly proportional to the inverse of the number of molecules observed on average in the confocal volume.

## OVERVIEW OF THE PROCEDURE

The protocol uses single color FCS to obtain one calibration curve for one type of FP (see Figure 2 for a schematic overview of the procedure). First, cells of interest are seeded and prepared for imaging (steps 1|-9|). Second, FCS measurements and fluorescence images are acquired to later determine a calibration curve (steps 10|-21|). Additional 3D confocal time-lapse movies can be acquired for further quantification (step 22|). This step does not require additional FCS measurements given that the same imaging settings are maintained. Data are processed to compute a calibration curve (steps 23|-29|). The images acquired in step 22| are converted to concentrations and protein numbers using the estimated calibration parameters (step 30|).

**Figure 2:**
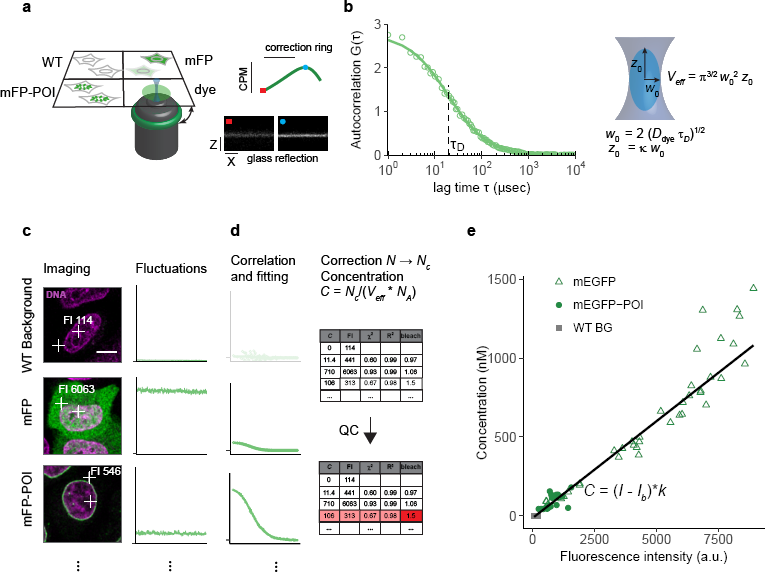
Workflow for FCS-calibrated imaging. (a) Setup of the multi-well chamber with four different samples and optimization of the water objective correction for glass-thickness and sample mounting using a fluorescent dye and the reflection of the cover glass. dye: fluorescent dye to estimate the effective confocal volume, WT: WT cells, mFP: WT cells expressing the monomeric form of the FP, mFP-POI: cell expressing the fluorescently labeled POI.
(b) Computation of parameters of the effective confocal volume using a fluorescent dye with known diffusion coefficient *D_dye_* and similar spectral properties as the fluorescent protein. Fitting of the ACF to a physical model of diffusion yields the diffusion time *τ_D_* and structural parameter *к.* The two parameters are used to compute the focal radius *w_0_* and the effective volume *V_eff_* (**Box 2**).
(c) Images and photon counts fluctuations are acquired for cells that do not express a fluorescent protein (WT), cells expressing the mFP alone (mEGFP), and cells expressing the tagged version of the POI, mFP-POI (mEGFP-NUP107). The image FI at the point of the FCS measurement is recorded. Scale bar 10 μm.
(d) The ACF is fitted to a physical model to yield the number of molecules in the effective volume. WT concentrations are set to 0. After background and bleach correction concentrations are computed using the *V_eff_* estimated in (a-b) and the Avogadro constant *N_A_*. Data is quality controlled with respect to the quality of the fit and the amount of photobleaching using objective parameters.
(e) Concentrations estimated from FCS and image fluorescence intensities are plotted against each other to obtain a calibration curve (black line).

For the sample preparation it is convenient to use a multi-well glass bottom chamber containing separate wells with cells expressing the fluorescently tagged POI (mFP-POI), cells expressing a monomeric form of the fluorescent tag alone (mFP), non-expressing wild-type (WT) cells, and a droplet of a reference fluorescent dye with a known diffusion coefficient. The fluorescent dye is used to adjust the sample and compute the effective confocal volume *V_eff_* (**Box 2**, steps 17|, 20| and 25|). Cells without a fluorescent tag are used to obtain the background photon counts for correcting the FCS derived concentrations as well as background image FI. Cells expressing the mFP alone are used to calibrate the monomeric fluorophore and estimate whether the POI oligomerizes. Furthermore, they provide a large dynamic range of concentrations and fluorescence intensities at different expression levels for estimation of the calibration parameters.

The objective as well as the bottom of the sample should be carefully cleaned and mounted to avoid any non-planarity between objective and cover-slip (steps 11 and 15|) as this leads to significant aberrations in the point spread function (PSF). Then the correction ring is adjusted and the fluorescent dye is measured (17|, **Box 2**). The microscope settings, such as pixel-dwell time, laser intensity, and detector gain, are carefully chosen for FCS and imaging of the mFP-POI (step 18| and 19|). The imaging settings must be kept the same for all images that need to be quantitatively analyzed and compared. The data for the calibration curve is obtained by placing 2-4 FCS measurement points at specific locations in the cell, typically at the two largest compartments, namely nucleus and cytoplasm (step 21|). We provide macros for the ZEN software to simplify the workflow and link the images with the FCS measurements (**Supplementary Software 1-3**). With *FCSRunner* (**Supplementary Software 1**) the user manually selects cells and FCS measurement points (step 21| Option A). The image and FCS measurements are then acquired with the previously determined imaging and FCS settings. Image pixel coordinates of the FCS measurements are automatically stored for further processing. This procedure is repeated for several cells and the three cell lines WT, mFP, and mFP-POI (Fig. 2c). The protocol also provides an optional fully automated workflow to yield reproducible measurements and a higher throughput (**Box 1**, step 21| Option B, **Supplementary Software 2** and **3**). Given that the laser intensity remains stable and that the imaging conditions are not changed, 3D stacks and 4D movies can now be acquired from cells in the multi-well plate without additional FCS calibration measurements (step 22|).

Background counts and the effective confocal volume are computed using *FCSFitM* (steps 23|-25|, **Supplementary Software 4**). For the fluorescent proteins ACFs, bleach and background correction factors are computed using *Fluctuation Analyzer 4G (FA)* ^18^ (step 26|). At the FCS measurement point the image fluorescence intensity *I* is calculated in a small region of interest (ROI) *(FCSImageBrowser*, **Supplementary Software 4**) (step 27|). The ACFs are then fitted to physical models of diffusion to obtain the number of molecules *N* (see **Supplementary Note 2**). A bleach and background corrected number of molecules *N_c_* is computed using the previously determined factors (see **Supplementary Note 3**) to yield a concentration *C* at the FCS measurement point

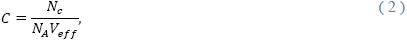

where *N_A_* is the Avogadro constant. The fitting and concentration estimation can be performed using *FCSFitM* (**Supplementary Software 4**) (step 28|). Finally, the data is quality controlled to correct for poor fits, measurements at the border of a compartment or on immobile structures (e.g. a nucleoporin on the nuclear envelope) (Fig. 2d and Supplementary Fig. 1) (steps 27|-28|). The concentrations are then plotted against the FI and fitted according to the following linear relationship (Fig. 2e):

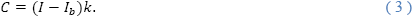

Here, *I_b_* is the mean background FI estimated from WT cells and *k* is the calibration factor. The last steps can be performed using the application *FCSCalibration* (step 29|, **Supplementary Software 4**).

The image pixel fluorescence intensities *I_p_* are converted to concentrations *C_p_* using Eq. (3) and to protein number per pixel *N_p_* (Fig. 3a) using the physical size of the 3D voxels, characterized by the sampling Δ*x* and Δ*y* and the Z-slice interval Δ*z*:

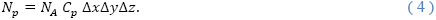

The conversion of fluorescence intensities is automatically performed using the ImageJ plugin *FCSCalibrate* (step 30|, **Supplementary Software 4**). See section "Quantification of FCS-calibrated images" for further details on quantifying protein numbers on cellular structures.

## APPLICATIONS OF THE PROTOCOL

The method described in this protocol is generic and can be used for different protein expression systems. Nevertheless, to achieve physiological expression levels, it is preferable to homozygously tag proteins at their endogenous genomic loci. The method is not restricted to cell monolayers, however, depending on the system, autofluorescence or scattering may impair reliable FCS measurements and imaging. The principles of this protocol are not restricted to the described microscopy setup and can be applied on all systems with photon counting detectors. We extensively tested and applied the method using mEGFP as green FP ^11–13,15,19^ and tested mCherry as a red FP. Other FPs can also be used as long as they are monomeric and have a fast maturation time (see ^20^ and http://www.fpvis.org). As alternative to mEGFP, mNeonGreen has been shown to be brighter and more photostable ^21^. The FP mCherry ^22^ is a good compromise of high photostability and fast maturation but has a rather low quantum yield. Depending on the application, other proteins, such as the recently described brighter mScarlet ^23^, may be better suited. Brighter and more photostable proteins could allow for longer imaging and improved signal to noise ratios.

This protocol, in combination with image analysis, is a powerful tool to develop quantitative models of cellular processes. For instance, this protocol can be used to estimate the amount of endogenously fluorescently labelled proteins in different cellular compartments throughout the cell cycle ^15^ (see **Supplementary Note 4**). Furthermore, this protocol can be applied to determine the stoichiometry and number of proteins in multi-protein complexes such as nuclear pore complexes ^24^, kinetochores ^25,26^, or centrosomes ^27^. Such structures should be resolved as single point sources by imaging ^14^ (see **Supplementary Note 5**). The density of proteins on membranes (Fig. 3, see **Supplementary Note 6**) or at the surface of macro-molecular complexes (e.g. chromosomes) ^13^ can also be estimated using this protocol. Finally, the number of proteins bound to a large macro-molecular complex can be monitored over time (e.g. condensins bound to chromatin ^12^).

**Figure 3:**
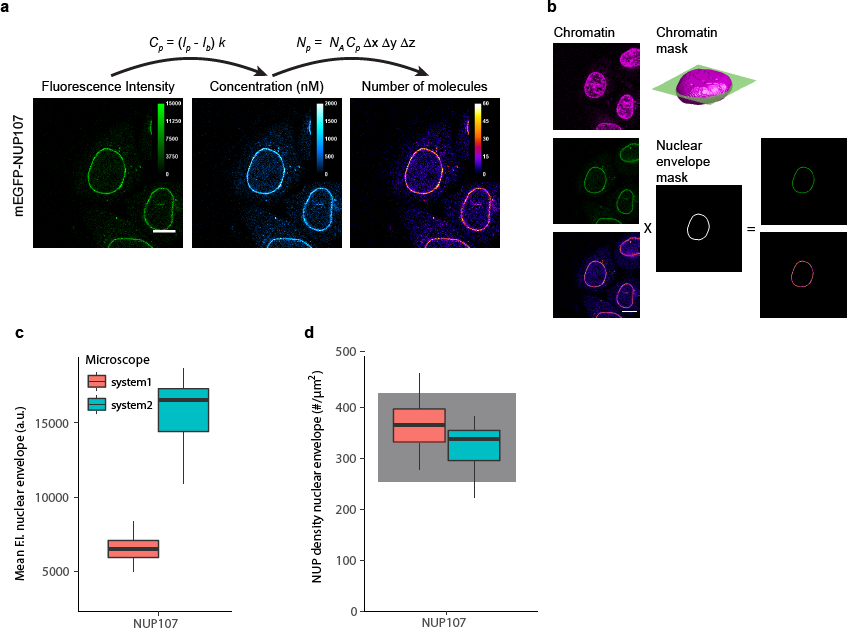
Example of image quantification using FCS calibration. (a) The FI at each pixel, *I_p_*, is converted to concentrations *C_p_* and protein numbers *N_p_* (for 3D and 4D images). The images are acquired with the same imaging parameters as the images used to compute the FCS calibration curve. Shown is data for a HeLa Kyoto cell line with endogenous NUP107 tagged N-terminally with mEGFP. Scale bar 10 μm.
(b) The quantitative distribution of a POI can be derived by using markers for cellular structures. DNA stained with SiR-DNA is used to compute a 3D chromatin mask of the nucleus. In the equatorial plane, a three pixel wide rim defines the nuclear envelope. Fluorescence intensities and protein numbers on the nuclear envelope can then be calculated. Scale bar 10 μm.
(c) mEGFP-NUP107 average FI on the nuclear envelope. Data shows results obtained on two different microscopes. The boxplots show median, interquartile range (IQR), and 1.5*IQR (whiskers), for *n* = 16-22 cells and 2 independent experiments on each microscope. System 1: LSM880, FCS and imaging using the 32 channel GaAsP detector. System2: LSM780, FCS using the APD detector (Confocor-3), imaging using the 32 channel GaAsP detector.
(d) Conversion of FI to protein numbers using the corresponding calibration curve obtained for each experiment and microscope system (not shown). The protein density on the nuclear envelope has been computed according to Eq. S24 (**Supplementary Note 6**). The gray shadowed boxes show the expected protein numbers for NUP107 ^18,19^.

## LIMITATIONS

This protocol is designed to quantify live cell images, since FCS measurements cannot be made in fixed cells. The amplitude of the autocorrelation function *G(τ)* decreases with the number of particles, thus the sensitivity of FCS decreases for high concentrations of the fluorophore. For a fluorophore such as EGFP, concentrations up to 1-2 μM can be measured. This limitation does not apply to the imaging of FI where the time integrated detector signal is used. In this case concentrations above the μM range can be measured as long as the detector is not saturated. At very low fluorophore concentrations, the method is impaired by scattering and autofluorescence. Nevertheless, by using appropriate corrections for the background, EGFP concentrations as low as a few pM can be measured (**Supplementary Note 3**). A dynamic range of pM to μM concentrations means that over 70% of the proteome of a human cell line is accessible to the presented method ^28^.

This protocol uses the fluorescence of the FP-tagged POI as a proxy for the protein distribution. It is important to test that the FP-tagged protein is functional (see also accompanying Nature Protocols ^4^). Furthermore, all the limitations of FPs such as potential dimerization induced by the FP, incomplete maturation, and photobleaching also apply to this protocol. Therefore fast maturing, monomeric and photostable FPs with a high quantum yield are preferable as tag. The protocol has been tested with mEGFP and mCherry, two FPs that have among the lowest tendency to dimerize, with dissociation constants above 70 mM^22,29,30^. From the nearly 30 fluorescently tagged proteins we investigated ^15^, the median whole cell concentration was 80 nM with a 90th percentile interval of [13-415] nM and a maximal value below 20 μM. This indicates that the number of expected dimers is negligible (maximally 1 dimer out of 10.000 proteins).

Photobleaching of the FP leads to underestimation of the number of proteins. In particular, point FCS with its long measurement times can cause strong photobleaching. To milder this issue and derive a reliable calibration curve, we suggest minimizing imaging and FCS time by acquiring one plane and measuring 1 min of FCS per cell. Imaging-induced bleaching was negligible with the described settings used for mEGFP, whereas the FCS-induced photobleaching, estimated by the decrease in photon counts, remained small (1-5%). For mCherry, photobleaching during FCS is stronger (5-15%), however, using the correction method as described in ^18^ and **Supplementary Note 3**, the effect of photobleaching can be accounted for. Therefore, with the settings used in this protocol. the effect of photobleaching in defining the calibration curve is negligible. For time-lapse 4D movies the bleaching strongly depends on the application and type of imaging. By imaging 40 time points with a time interval of 90 sec and 31 planes per time point (0.25×0.25×0.75 μm pixel resolution), we have not observed a significant photobleaching for POIs tagged with mEGFP ^15^. Higher frame rates, laser power or increased spatial resolution may cause stronger photobleaching which needs to be accounted for. To do this, in a first step, fluorescence intensities are converted to concentrations using the calibration curve. In a second step the calibrated images are scaled using an estimated bleaching function ^31,32^.

In order to become fluorescent, FPs must undergo a maturation step that can last minutes to hours. The proteins we tested in this protocol, namely mEGFP and mCherry, have reported maturation times from 10-60 min ^22,23,33–35^. This delay leads to an underestimation of the number of proteins, whereby its extent depends on the half-life of the POI. The fraction of mature FP with respect to the total protein amount can be approximated by *k_M_/(k_d_ + k_M_)* (ref. ^35^), where *k_M_* and *k_d_* are the rate constants for maturation and degradation, respectively. The half-life of proteins (∼1 /*k_d_*) varies from over 1 h to several 10 of hours (median 46 h, 90th percentile [8-184] h; ^36^). Therefore, by assuming a maturation time of 0.5 h, for 90% of all human proteins the error due to delayed maturation ranges from 0.25% to maximally 6% of the total protein amount.

This protocol is not restricted to the use of mEGFP or mCherry. Depending on the experimental system, issues with photobleaching or maturation time can also be addressed by using more stable and brighter proteins such as mNeonGreen or mScarlet ^21,23^.

## COMPARISON TO OTHER METHODS

In the past, several approaches have been developed to obtain quantitative readouts from fluorescent images (reviewed by ^37,38^). With the possibility to express fluorescently tagged proteins from their endogenous locus, these methods can now be used for proteomics studies in almost any eukaryotic cell. Typically, quantitative microscopy relies on the computation of a calibration function to convert fluorescence intensities to physical quantities. The calibration function can be computed using intra- or extracellular fluorescent standards ^39,40^, or, as we present in this protocol, direct measurements of the concentrations within cells ^41–43^. Protein numbers can also be directly measured by step-wise photobleaching ^44,45^, photon emission statistics ^45–47^, or, in compartments with freely diffusing components, by analyzing fluorescence fluctuations ^5,6^

Extracellular fluorescent standards require purified FP, the same FP as used for tagging the POI. Depending on the application, solutions of known concentrations ^48^ or diffraction-limited complexes with known stoichiometry (e.g. Virus-like particles ^49^) are measured along with the cells expressing the FP-tagged POI. The method is relatively simple but requires additional sample preparation, knowledge of stoichiometry and concentration, and most importantly does not ensure whether the intra- and extracellular excitation and emission properties of the FP are the same ^38^. An alternative is the measurement of total protein amounts with quantitative immunoblotting ^37,39^ or mass spectrometry ^28,36^. Using additional specialized equipment and reagents, both methods provide an estimate of the total protein amount that can be related to the total cell fluorescence. However, measurements are taken at the population level and the total amount of protein per cell needs to be extrapolated. Inaccuracies arise in the conversion of immunoreactivities to protein amount, estimate of total number of cells per immunoblot, and when the total protein level strongly differs between cells (e.g. cell cycle variations).

Intracellular calibration standards circumvent the issue of differences in fluorescence in an extracellular environment but require an a priori knowledge of the stoichiometry or concentration. This method has been extensively used in yeast with proteins on the kinetochore as calibration standard ^40,48,50^. In mammalian cells an intracellular calibration standard has not been agreed on, making the method not yet applicable.

The direct measurement of protein numbers using discrete photobleaching events is not suited for long time imaging. However, it can be used to define intracellular calibration standards ^45^. Counting by photon statistics requires a specialized hardware with at least four detectors and bright fluorescent probes ^46,47^ The possibility to tag proteins with organic dyes in live cells using endogenous expression of proteins tagged with SNAP ^51^, CLIP ^52^ or Halo ^53^ could make the method more accessible in the future.

In this protocol the calibration curve is computed from FCS concentration measurements in cells. Compared to other methods that use calibration standards, this method requires a specialized confocal setup that can then be used for further imaging. Data analysis requires parameter estimation and fitting. Thanks to the availability of software solutions (e.g. QuickFit3 http://www.dkfz.de/Macromol/quickfit; SimFCS, https://www.lfd.uci.edu/globals/; **Supplementary Software 4** in this protocol) the user can perform the analysis without a specialized knowledge of FCS.

## EXPERIMENTAL DESIGN

### Cellular samples

For FCS-calibrated imaging three different cell line samples are used. To measure the background autofluorescence and background photon counts the cell line of interest is needed as a WT clone, i.e. not expressing recombinant FPs. Alternatively, measurements in the culture media can be taken. However, this typically underestimates the background by about 10%. To obtain concentration and FI measurements that span a range of up to 2 μM, cells expressing the FP alone are needed. These cells can be generated by transiently transfecting WT cells with a plasmid containing the monomeric form of the FP used for tagging of the POI. To achieve physiological expression of the POI, genome edited cells (ZFN, CRISPR/Cas9) expressing the POI tagged with the FP are preferred (see our accompanying Nature Protocol ^4^). The criteria for FPs in FCS are the same as for fluorescent imaging: (*i*) high molecular brightness, (*ii*) fast maturation time, (*iii*) minimal oligomerization, (*iv*) high photostability, (*v*) spectral match with available laser lines and filters. Monomeric EGFP and monomeric EYFP or other bright green/yellow FPs are well suited for FCS-calibrated imaging ^21^. The FP mCherry has been tested and is suitable as red FP. Other monomeric and potentially brighter and more photostable red fluorescent proteins can also be used ^23^.

### FCS settings

The laser intensity should be sufficiently high to obtain good counts per molecule (CPM), as this will lead to a good signal to noise ratio, but also sufficiently low to avoid photobleaching, photodamage and saturation effects during the FCS measurement. First set the FI for the biological sample expressing the FP only to obtain CPMs from 1 to 4 kHz with low photobleaching. In point FCS, photobleaching can be assessed by a continuous decrease in photon counts. Measuring few FCS points (2-4) per cell in compartments where the POI is freely diffusible minimizes photobleaching. Furthermore, using *FA*, ACFs and protein numbers corrected for photobleaching can be computed (see **Supplementary Note 3**). The choice of the pinhole size is a tradeoff between detected signal, sensitivity, and the quality of the fits. Larger pinholes cause a decrease in CPM leading to a decrease in the quality of the fit. This lowers the precision of the diffusion time estimate, a parameter affecting the precision of the effective confocal volume estimate. We found that pinhole sizes from 1 to 1.6 Airy Units (AU) gave reliable confocal volume estimates without impairing the quality of the fits (Supplementary Fig. 2a and ^54^). Since the size of the effective confocal volume depends on the laser power and pinhole size (Supplementary Fig. 2b-c), the same parameters used for the FCS measurements of the POI must also be used for the reference dye.

### Imaging settings

We distinguish between images that are used to compute the calibration curve and images where we apply the image calibration. To define the parameters of the calibration curve, single plane images at single time-points are best suited. Saturation of the image should be avoided, as this will preclude quantification above an upper limit. Furthermore, photobleaching during the acquisition of the calibration images should be minimal. This is typically the case as only one single image per cell is required. To compute an approximation of the protein numbers in a measurement point, the method uses the 3D pixel dimensions of the imaging system. Consequently, images that need to be quantified must be 3D (XYZ) or 4D (XYZT) image stacks. In time-lapse movies, the effect of photobleaching ^31,32^ can affect the quantification. Photobleaching can be reduced by using mild imaging conditions (e.g. shorter pixel dwell-times, less Z-planes) or more photo-stable FPs ^21,23^. It is recommended, to estimate the photobleaching parameters by fitting a function (e.g. exponential) to the total intensity changes of repeatedly acquired images. These parameters can then be used to correct the whole 4D time-lapse after calibration. The spatial sampling must be high enough in order to resolve the structure of interest. For structures that are close to the size of the point spread function (PSF) or non-isotropic, such as membranes, a spatial sampling close to the Nyquist criterion should be used. Similarly to the FCS measurement, the imaging pinhole needs to be adjusted and should be small enough for good confocal sectioning (also 1-1.6 AU). If possible use the same pinhole for imaging and FCS.

### Quantification of FCS-calibrated images

The calibrated images allow an estimate of the protein distribution in specific cellular compartments, organelles or structures. For that purpose additional fluorescent reference markers of the structure of interest imaged in an independent fluorescent channel are required (see also “Markers to aid image segmentation”). For instance, to separate proteins on the nuclear envelope from proteins in the nucleus and cytoplasm a nuclear or DNA marker can be used. Quantification of total protein numbers on structures that are within the size of the PSF, such as large multiprotein complexes or membranes requires particular care ^13,14^ As shown in the supplement the fluorescent signal needs to be integrated throughout the structure in 3D (Supplementary Fig. 3). A diffraction-limited structure requires integration of the signal in a 3D volume of the size of the PSF (Supplementary Fig. 3c-d). If the size of the PSF is known, one can approximate the total number of proteins from a subset of pixels (Supplementary Fig. 3c-d, squares and diamonds). Furthermore, spatial symmetry properties of the structure of interest can be used. For example, in the equatorial plane of the nucleus, the nuclear membrane can be assumed to be locally isotropic in Z. In this case, to obtain the density of proteins on the nuclear membrane, it is enough to estimate the protein number per unit length on a sufficiently broad nuclear rim at the equatorial plane (Fig. 3 and Supplementary Fig. 3f).

### Markers to aid image segmentation

To quantify the total number of molecules in the whole cell or in specific subcellular compartments or structures, fluorescent markers and image processing are required to define discrete volumes. Cellular compartments useful to be stained for image segmentation and further quantification are the nucleus/chromatin and the cytoplasm. The cytoplasm can be obtained from the difference between a segmented whole cell volume and the segmented nucleus/chromatin compartment. Note that fluorescent markers enabling cellular or subcellular image segmentation must not interfere/cross-talk with the FP-tagged to the POI.

DNA counterstains suitable for live cell imaging are SiR-DNA ^55^ and Hoechst 33342. SiR-DNA has a low cytotoxicity and can be excited with a 633 nm laser allowing for long-term imaging. Instead of a chemical stain, fluorescently labeled histone 2B (H2B) can be used ^56^. An easy solution to segment the cell surface is to use fluorophores that stain the extracellular culture media and cannot penetrate the cellular membrane ^12,15^, such as fluorophores linked to a bulky molecule (e.g. high molecular weight dextran or IgG). This negative stain of the cell boundary has the advantage of being easily segmentable and the fluorescent dye can be adapted to the combination of DNA stain and protein of interest fluorescent probe. For instance, a suitable dye in combination with mEGFP and SiR-DNA is DY481XL. This large stokes-shift dye can be excited together with mEGFP at 488 nm but emits in the red spectral range, this reduces the need for an additional excitation laser and decreases phototoxicity. A list of fluorophores that work well in combination with the commonly used FPs mEGFP and mCherry are provided in Table 1.

**Table 1:** Combination of fluorescent proteins and dyes that can be used for tagging the POI and staining cellular landmarks.

| Fluorescent protein for POI | Dye/fluorescently tagged protein or nuclear landmark | Dye for cell boundary landmark | Remarks |
| --- | --- | --- | --- |
| *mEGFP*<br>Ex: 488<br>Em: 492-544 | *SiR-DNA*<br>Ex: 633<br>Em: 640-695 | *Dy481XL-01*<br>Ex: 488<br>Em: 640-695 |  |
| *mCherry*<br>Ex: 561<br>Em: 588-690 | *H2B-mCerulean*<br>Ex: 458<br>Em: 458-492 | *Atto430LS-31*<br>Ex: 458<br>Em: 588-690 | Small bleed-through of mCerulean in the Atto430LS-31 channel |
| *mEGFP*<br>Ex: 488<br>Em: 492-544 | *H2B-mCherry*<br>Ex: 488 or 561<br>Em: 588-624 | *Dy481XL-01*<br>Ex: 488<br>Em: 625-695 | Cross-talk between mCherry and Dy481XL-01 channel |
| *mCherry*<br>Ex: 561<br>Em: 588-690 | *Hoechst 33342*<br>Ex: 405<br>Em: 410 - 480 | *Alexa488*<br>Ex: 488<br>Em: 492-544 | Phototoxicity of 405 nm laser upon long-term imaging |
| *mCherry*<br>Ex: 561<br>Em: 588-610<br>(small emission range to minimize bleed-through) | *SiR-DNA*<br>Ex: 633<br>Em: 640-695 | *Atto430LS-31*<br>Ex: 458<br>Em: 588-690<br>or *Alexa488*<br>Ex: 488<br>Em: 492-544 | Bleed-through of SiR-DNA in the mCherry channel when excited with 561 nm. This impairs the FCS measurements at low protein numbers |

The first 3 entries represent optimal combinations. For low expressing proteins the chromatin stain should not cause crosstalk into the FP channel. The emission range is given as a guideline for minimizing crosstalk. Ex: Excitation, Em: Emission.

### Controls needed

A serial dilution of the fluorescent dye is used test the linearity of the imaging detector and the FCS concentration measurements. The concentration of the fluorescent dye can be estimated by measuring its absorption using the dye extinction coefficient as provided by the manufacturer. To test the linearity of the imaging detector, measure the different dilutions with the settings used for imaging the POI and plot the average intensity against the dilution factor. Similarly, perform FCS measurements and compute the concentrations (**Supplementary Notes 2** and **3**). A linear relationship between computed concentrations and dilution factor is expected for low dye concentrations, whereas at high dye concentrations one observes a deviation from linearity. In our setup, and with Alexa488/568, the limit was approximately reached at 2 μM. The linearity range for the imaging detector is larger than for the FCS measurements. This allows estimating the calibration parameters from cells with a low (<1-2 μM) FP concentration and apply the calibration coefficients to cells with higher FP concentrations. In the calibration curve obtained from *FCSCalibration* (**Supplementary Software 4**), deviation from linearity is easily detected by visual inspection. In order to remove cells with a high FP concentration and improve the quality of the calibration curve an upper limit for the concentration can be set in *FCSCalibration.* To test the stability of the microscopy calibration, repeated FCS and imaging data are acquired over a longer period (e.g. 12 h). Such long-term experiments can be performed using the tools for the automated imaging (**Supplementary Software 2** and **3**). In our setup, we did not observe systematic changes in the calibration function or CPMs over a period of 12 h.

To identify POI oligomerization, a cell line transiently expressing the mFP alone is used. A larger mean CPM in the POI cell line compared to the mean CPM measured in the mFP cell line indicates oligomerization. Inspection of the CPM distribution can be used to identify distinct populations. In *FCSCalibration,* oligomerization is considered by computing a correction factor from the ratio of the two POI CPM to mFP CPM. Cell autofluorescence, can be accounted for by measuring cells that do not express the FP. Finally, to test the precision of the image quantification obtained from this protocol, the user can use a priori knowledge on specific POIs. For instance, in this protocol we used nucleoporins from which we know the nuclear pore complex (NPC) density and stoichiometry in HeLa cells. Alternatively, the estimated whole cell protein numbers obtained with this protocol can be compared to results obtained from other quantitative methods (see **Comparison to other methods**).

## MATERIALS

## REAGENTS

### Cell lines

#### CAUTION

Cell lines used in your research should be regularly checked to ensure they are authentic and not infected with mycoplasma.

- Cell line with a FP labeled POI. Preferably use an endogenously tagged cell line. Here we show data for a HeLa Kyoto cell line with mEGFP-NUP107 (endogenously tagged using a zinc finger nuclease) ^57^ The cell line is available upon request.

#### CRITICAL

The FP must be appropriate for the available laser lines and filters. This protocol has been tested with mEGFP and mCherry. Other monomeric fluorescent proteins can also be used.

- Cell line of interest as WT clone. Here we used a HeLa Kyoto cell line. The HeLa Kyoto cells can be obtained from Dr. S. Narumiya, Department of Pharmacology, Kyoto University.

### Mammalian cell culture

- High Glucose Dulbecco’s Modified Eagle Medium (DMEM) (Life Technologies, cat. # 41965-039).
- Fetal Bovine Serum (FBS, Life Technologies, cat. # 10270106).
- 10000 units/ml Penicillin–Streptomycin (Life Technologies, cat. # 15140122).
- 200 mM L-Glutamine (Life Technologies, cat. # 25030081).
- NaCl (Sigma Aldrich, cat. # S5886).
- KCl (Sigma Aldrich, cat. # P5405).
- Na_2_HPO_4_ (Sigma Aldrich, cat. # 255793).
- KH_2_PO_4_ (Sigma Aldrich, cat. # S5136).
- HCl (Sigma Aldrich, cat. # H1758).

#### CAUTION

HCl is highly corrosive and should be used with the appropriate protective equipment.

- 0.05% (vol/vol) Trypsin (Life Technologies, cat. # 25300054).

### Live cell imaging

- Cell culture buffers suitable for imaging. These should be non-fluorescent and have a suitable refractive index. For the protocol presented here, we used CO_2_-independent imaging medium without phenol red (Gibco, cat. # ME080051L1).

### Confocal volume calibration

- Bright and photostable dyes that match the spectral properties of the FP with known diffusion coefficients are required. For example Alexa488 (Thermo Fisher, NHS ester, cat. # A20000) or Atto488 (Atto-Tec, NHS ester, cat. # AD 488-31) to match mEGFP and Alexa568 (Thermo Fisher, NHS ester, cat. # A20003) to match mCherry.

### Transfection

- Plasmid expressing the FP used for tagging the POI under a mammalian promoter. Here we used pmEGFP-C1 kindly provided by J. Lippincott-Schwartz (Addgene plasmid # 54759). mEGFP is the monomeric form of EGFP with the A206K mutation.
- Opti-MEM® I Reduced Serum Medium, GlutaMAX™ Supplement (Life Technologies, cat. # 51985-026).
- Fugene6 (Promega, cat. # E2691).

### Cellular markers (optional)

- SiR-DNA^55^, also known as SiR-Hoechst (Spirochrome, cat. # SC007).
- Hoechst-33342 (Thermo Fisher, cat. # 62249).

#### CAUTION

SiR-DNA and Hoechst-33342 are mutagenic and harmful if swallowed. It causes skin and respiratory irritation. It is suspected of causing genetic defects; handle it while wearing appropriate personal protective equipment. Keep it protected from light.

- Dextran, Amino, 500,000 MW (Life Technologies, cat. # D-7144).
- NaHCO_3_ (Sigma Aldrich, cat. # S576).
- Dy481XL-NHS-Ester (Dyomics, cat. # 481XL-01).
- Atto430LS-31 NHS Ester (Molecular Probes, cat. # AD-430LS-31).
- Molecular Probes® DMSO (Thermo Fisher, cat. # D12345).

### Other reagents

- Pure ethanol for cleaning the objective (Sigma Aldrich, cat. # 1009831000).

## EQUIPMENT

### Hardware requirements

- Scanning confocal microscope with fluorescence correlation setup. The detectors for fluorescence correlation must work in photon counting mode. For imaging the detectors must be in a linear response range (see EQUIPMENT SETUP).
- Temperature control chamber for long time imaging of living cells.
- A high numerical aperture water-immersion objective. On Zeiss systems use C-apochromat Zeiss UV-VIS-IR 40x 1.2 NA, specially selected for FCS (421767-9971-711).
- Observation chamber with coverglass bottom suitable for cell culture with at least 4 separate wells. Imaging plates 96CG, glass bottom (zell-kontakt, cat. # 5241-20), 4 to 8 well IBIDI (IBIDI, cat. # 80427, 80827), 4 to 8 well LabTek (Thermo Fisher, cat. # 155382, 155409, 155383, 155411) both #1 or #1.5 glass thickness can be used.
- For long time imaging an objective immersion micro dispenser system should be used. Leica provides a commercial water immersion micro dispenser.
- 15 cm Nunc dishes for cell culture (Sigma-Aldrich, cat. # D9054).
- PD-10 Sephadex desalting columns (optional, GE Healthcare, cat. # 52130800).
- Slide A-Lyzer 10.000 MWCO (optional, Thermo Fisher, cat. # 66810).
- Viva spin 30.000 MWCO (optional, Sigma-Aldrich, cat. # Z614041-25EA).
- 0.22 μm Filters (Sigma-Aldrich, cat. # SLGP033RS).
- Whatman™ lens cleaning tissue, grade 105, 100×150 mm (GE Healthcare Life Sciences; cat. # 2105-841).
- Silicon grease for sealing (KORASILON™ paste, highly viscous Obermeier; cat.# 8000054-99).
- TetraSpeks microspheres, 100 nm (optional, Thermo Fischer, cat. # T7279).
- Workstation with Windows 7 or higher.

### Software requirements

#### CRITICAL

All custom software packages are provided as **Supplementary Software 1–4**. An overview of software requirements is provided in **Supplementary Table 1** with links to the most recent source.

- Microscope software licensing for FCS data acquisition.
- For Zeiss microscopes, ZEN black edition (version higher or equal to ZEN 2010, https://www.zeiss.com/microscopy/us/downloads/zen.html) to run the VBA macros (**Supplementary Software 1** and **2**).
- FiJi (https://fiji.sc) ^58,59^ installation on analysis computer.
- For high-throughput FCS and imaging data acquisition with adaptive feedback (21| Option B), FiJi installation on the computer with the ZEN software controlling the microscope and **Suplementary Softare 3**.
- *Fluctuation Analyzer 4G (FA, ^18^, https://www-ellenberg.embl.de/resources/data-analysis)* on the analysis computer to compute corrected autocorrelation traces from the raw photon counting data.
- The workflow for FCS fitting (*FCSFitM*, **Supplementary Software 4**) is a MATLAB tool compiled for Windows. This does not require a MATLAB installation. To run the source code on a different operating system, MATLAB with the toolboxes optimization and statistics needs to be installed (R2014a or later, The MathWorks, Inc., Natick, Massachusetts, United States).
- The data analysis tool *FCSCalibration* (**Supplementary Software 4**) is a *R* tool *(http://www.R-project.org)* with a *shiny* graphical user interface *(http://cran.r-project.org/package=shiny).* We recommend to use the source code in combination with *RStudio (https://www.rstudio.com).* In the packaged executable version for Windows all required dependencies are installed.

## REAGENTS SETUP

### HeLa growth medium

High Glucose DMEM supplemented with 10 % (vol/vol) FBS, 100 U/ml Penicillin, 100 μg/ml Streptomycin and 2 mM L-Glutamine. Supplemented DMEM medium is abbreviated as complete DMEM. The growth media can be stored up to 1 month at + 4 °C. Pre-warm the solution in a + 37 °C water bath before use.

### Imaging medium

CO_2_ independent medium without phenol red supplemented with 10% (vol/vol) FBS, 2 mM L-Glutamine, 1 mM sodium pyruvate. Eventually add glucose to achieve a final glucose concentration as in the complete DMEM (4.5 g/liter). Filter through a 0.22 μm membrane to clear the medium of precipitants. The imaging media can be stored up to 1 month at + 4 °C. Pre-warm the solution in a + 37 °C water bath before use.

### PBS, 1X

Prepare 137 mM NaCl, 2.7 mM KCl, 8.1 mM Na_2_HPO_4_ and 1.47 mM KH_2_PO_4_ in ddH_2_O. Adjust the pH to 7.4 with HCl and autoclave. The solution can last several months.

### SiR-DNA

Dissolve the SiR-DNA in DMSO and store at - 20 °C according to manufacturer’s instructions. The solution can be thawed multiple times and last several months. Aliquoting is not recommended.

### Fluorescent dyes for calibration

Dissolve the fluorescent dye in DMSO according to manufacturer’s instructions. Dilute a small volume of the stock solution in ddH2O to the desired concentration (5-50 nM). Store the dye solutions at + 4 °C. The solution lasts for several months.

### Dextran labeled with a fluorescent dye

The protocol to generate dextran labeled with a fluorescent dye is given in **Box 3**. The reagent can be stored for > 1 year at - 20 °C.

## EQUIPMENT SETUP

### Detectors

For FCS on Zeiss LSM confocal systems GaAsP or APD detectors (Confocor-3 system or in combination with a PicoQuant upgrade kit) can be used. For Leica microscopes the HyD SMD detectors or APD detectors can be used. The protocol and software is based on Zeiss LSM microscopes (LSM780, LSM880) using ZEN black edition and an inverted Axio Observer. In particular, the VBA macros only work with ZEN black edition. For imaging we tested GaAsP detectors (LSM780 and LSM880) and the AiryScan detector (LSM880).

### Microscope objective

Use a water objective with a high numerical aperture and the best chromatic correction as recommended for FCS. Prior to objective purchase we recommend verifying the PSF and confocal volume, using fluorescent dyes (e.g. Alexa488 and Alexa568) and/or fluorescent beads. Use pure water (ddH2O) as immersion media as this yields the best optical properties. Oil immersion media that match the refractive index of water only do this at a specific temperature (typically 23 °C). Thus due to refractive index mismatch the results with oil immersion media may not have the same quality as the results obtained with water as immersion media.

### Supplementary Software Setup

The analysis and acquisition software is listed in Supplementary Table 1. For the installation of the custom packages (**Supplementary Software 1-4**) follow the instructions included in the software documentation. Most recent versions of the **Supplementary Software 1-4** can be found in *https://git.embl.de/grp-ellenberg/fcsrunner*, *https://git.embl.de/grp-ellenberg/mypic*, *https://git.embl.de/grp-ellenberg/adaptive_feedback_mic_fiji*, *https://git.embl.de/grp-ellenberg/FCSAnalyze* respectively. User guides can also be found on the respective Wiki pages, e.g. *https://git.embl.de/grp-ellenberg/fcsrunner/wikis*. Install all required software prior to starting the experiment.

### Installing Fluctuation Analyzer 4G (FA)

Download the software from *https://www-ellenberg.embl.de/resources/data-analysis* ^18^. On the data analysis computer with a Windows operating system unpack the software and run the setup.

### Installing FiJi

Download the latest version of FiJi from https://fiji.sc and install the software on the data analysis computer. For the adaptive feedback pipeline install the software on the computer that runs the software controlling the microscope.

### Installing R (optional)

In case *FCSCalibration* packaged for Windows is not used or does not work (**Supplementary software 4**), you need to install the *R* software. Download *R* (https://cran.r-project.org) or preferably *RStudio (https://www.rstudio.com)* and install the software on the data analysis computer.

## PROCEDURE

### Sample preparation TIMING 1 h hands-on, 1 d waiting time, 15 min hands-on

#### CRITICAL

To avoid contamination all steps should be performed in a laminar flow hood.

1| For imaging and FCS, a cell density of 50-70% confluence is recommended. Seed the cells on a glass bottom multi-well plate one day before the experiment. Prior to seeding, wash each well twice with PBS. For an 8 well LabTek chamber seed 1.6-2.0×10^4^ HeLa Kyoto cells into 300 μl complete DMEM per well. Seed cells expressing the fluorescently tagged POI in one well and WT cells in two separate wells. One well should be left empty for the reference FCS measurement with a fluorescent dye.
2| Grow cells in complete DMEM at 37 °C, 5% CO_2_. Wait for cells to attach (6 h) before transfection in Step 3.
3| Transfect the HeLa Kyoto cells with a plasmid containing the mFP of interest 6 h after seeding. Transfection can be carried out using FuGENE reagent. Mix 50 μl Opti-MEM with 1 μl FuGENE6 in an Eppendorf tube. Incubate for 5 min at room temperature (RT, 23 °C). Add 300 ng of plasmid DNA (we use pmEGFP-C1 as an example) and mix well. Incubate the transfection mixture for 15 min at RT. Add 25-50 μl of transfection complex to one well with seeded WT cells. Incubate the cells at 37 °C, 5% CO_2_ overnight. **CAUTION** Depending on the cell line transfection in the presence of antibiotics may impair cell health and transfection efficiency. If necessary, transfection can be performed in the absence of antibiotics. **CRITICAL STEP** Carefully dispense the transfection mix to avoid detaching of cells or contamination of neighboring wells. The amount of cDNA, transfection reagent or the time between transfection and imaging needs to be adapted. The goal is to achieve rather low expression levels of the mFP close to the expression levels of the mFP-POI.
4| On the next day, before preparing the sample for imaging (steps 5| to 9|), turn on the microscope, microscope incubator and all required lasers. Allow the confocal microscope system and the incubation chamber to equilibrate and stabilize. Depending on the setup, this can take over one hour.
5| Prior to imaging, change the medium to a phenol red free imaging medium. Use a CO_2_ independent medium if your microscope incubator does not allow for CO_2_ perfusion or if the CO_2_ perfusion impairs the FCS measurements.
6| For the automated acquisition of FCS data and imaging data (step 21| Option B), staining of the DNA is required. In order to do this: add SiR-DNA to the cells to achieve a final concentration of 50-100 nM. Incubate for 2 h at 37 °C (without CO_2_) until incorporation of SiR-DNA is complete. During the incubation time proceed with steps 7| to 20|. **CRITICAL STEP** In combination with a POI tagged with mCherry, the vital dye Hoechst 33342 can be used instead of SiR-DNA. To minimize toxicity, incubate cells for 10 min with 10 nM Hoechst and replace with fresh medium. The staining and cytotoxicity of the DNA dye can differ between cell lines and imaging protocols. This should be carefully tested for example by monitoring the mitotic timing.
7| (Optional) If required, add any further fluorescent markers for the cell boundary (see Table 1). **CRITICAL STEP** The presence of fluorescently labeled dextran in the cell culture medium reduces the staining of DNA by SiR-DNA. Cell toxicity and optimal SiR-DNA concentration should be tested in combination with the fluorescent marker in the cell culture medium.
8| Place a 20-50 μl drop of the diluted fluorescent reference dye (5-50 nM) in the empty chamber well. **CRITICAL STEP** This fluorescent dye should have similar spectral properties to the FP of interest and with a known diffusion coefficient. In this example we use Alexa 488.
9| To minimize liquid evaporation during the measurements, seal the multi-well plate chamber with silicon grease or use a gas permeable foil suitable for cell culture when using CO_2_ dependent imaging media. This step is recommended in case the humidity level is not sufficiently maintained in the microscope incubation chamber during data acquisition.

### Microscope setup and effective confocal volume measurement TIMING 1-2 h

10| Ensure that the confocal microscope system and the incubation chamber are equilibrated and stabilized (Step 4|). Select the beam path and emission filter/range for FCS and sample imaging, achieving the highest match between both. Select the emission filters to get highest emission signal but smallest (in the ideal case no) crosstalk between the individual fluorescence channels. Additional fluorescent channels may be needed for cellular markers (Table 1). The settings for the green (mEGFP) and red (mCherry) FPs as well as the fluorescent dyes Alexa488/568 are shown in Table 2. The emission range can be increased if only one fluorescent marker is present. Note that the protocol uses single color FCS to derive a calibration curve for a FP.
11| Clean the water-immersion objective and the glass bottom plate with lens cleaning tissue rinsed with pure ethanol. **CAUTION** The objective can be damaged if pressure is applied to the lens.
12| Make a rough adjustment of the water objective correction collar to match the glass thickness based on manufacturer's specification (#1 corresponds to 130-160 um, #1.5 corresponds to 160-190 μm).
13| Place a drop of pure water (ddH_2_O) on the objective and mount the sample.
14| Check correct and planar mounting of the sample. Planarity can be tested using the glass reflection in a XZ scanning mode, with the FCS objective at the smallest zoom or with a lower magnification objective. One should observe a horizontal straight line with negligible deviation in Z (<0.1 degrees tilting) (Fig. 2a). If the reflection of the glass is not planar, adjust your sample or the sample holder accordingly.
15| Wait for the system to equilibrate (∼30 min).
16| Meanwhile, create the folder structure to save the calibration data. ~~~
Your_Experiment_Folder\
     Calibration\
          dye\ contains the FCS measurements of the fluorescent dye
          WT\ contains the images and FCS measurements of the WT cells
          mFP\ contains the images and FCS measurements of the cells expressing the monomeric FP
          POI\ contains the images and FCS measurements of the cells expressing the POI tagged with the FP
~~~
17| *Pinhole, beam collimator, and objective correction collar adjustments.* Focus the objective on the well with the fluorescent reference dye. Set the pinhole size to 1-1.6 AU. Press the count rate tab. In the ZEN software the count rate window displays average photon counts and average CPM. The latter is calculated from the photon counts divided by an estimated number of molecules. Find the interface between the glass and the fluorescent solution. This is the *Z* position where the count rate and the total photon count suddenly increase. Move 30 μm above the glass surface inside the fluorescent solution. Select the laser power to achieve a CPM above 3 kHz and below approx. 30 kHz. An upper limit of the laser power is reached when the CPM does not increase with increasing laser power. Set the laser power well below (at least 1/2) this upper limit. The final laser power will be set using the FP as reference (step 18|). Close the count rate window. When using Zeiss LSM systems, perform both coarse and fine pinhole adjustments in X and Y for the channel of interest. Press the count rate tab and turn the correction collar to achieve the maximal count rate. Verify that this position corresponds to a maximal CPM.
18| *Setting FCS laser power using the mFP-POI.* Focus the objective in the well containing the cells expressing the FP-tagged POI. Acquire an image of the FP. Place an FCS measurement point in a region of the cell where the protein is expected to be freely diffusing, homogeneously distributed and free of aggregates (e.g. fluorescent aggregates in juxtanuclear regions) and has a low concentration. When cells exhibit variable expression levels, it is recommended to measure cells with a low expression level of the mFP-POI. Press the count rate tab and change the laser power to achieve a 4 kHz ≥ CPM ≥ 1 kHz whereby the total counts should not exceed 1000 kHz. Perform a FCS measurement over 30 sec to assess the photobleaching. Photobleaching can be assessed by a decrease in photon counts over time. **CAUTION** The APD and GaAsP detectors are very sensitive and can be damaged if too much light is used. Be careful not to exceed the maximal detection range. Stop acquisition if the detector shutter closes. **? TROUBLESHOOTING**
19| *Setting imaging conditions using the POI.* These imaging settings will be used to compute the FCS calibration parameters (step 29|). Set the pinhole size and laser power as close as possible to the values used for the FCS measurements (steps 17|- 18|). Set the pixel size, pixel dwell, and the zoom to be suitable and optimal for your sample. Set the detector gain value so that saturation is avoided. Saturation can be tested by analyzing the intensity histogram or visualized using the range indicator. Set the imaging conditions to acquire a single plane (2D images and no time-lapse). This is the image used to derive the FCS calibration parameters. For 3D imaging (XYZ stacks) use an uneven number of sections in order to always associate the central Z plane with the FCS measurements. Save the settings as a user-defined configuration in ZEN. **CRITICAL STEP** The pinhole size should be adjusted to achieve a good confocal sectioning for the structure of interest. The imaging settings determined in this step should be kept for further 3D or 4D imaging (step 22|) and calculating concentrations and protein numbers using the FCS calibration curve. **? TROUBLESHOOTING**
20| *Effective confocal volume measurement.* Move to the well containing the fluorescent reference dye. Focus 30 μm above the glass surface inside the fluorescent solution. Place the FCS measurement point in the center of the field of view. Use FCS settings identical to the settings for the POI (step 18|). Set the measurement time to 30 sec and 6 repetitions. Start the measurement. Save the data with the raw data for further processing to the *Calibration/dye* folder. **CRITICAL STEP** It is essential that the laser power and pinhole size are the same as determined in 18|. *FA* computes a bleach-corrected ACF using the raw counting data. Thus it needs to be ensured that this data is saved. For Zeiss LSM select in the Maintain/ConfoCor options `save raw data during measurement`. In the saving directory you should find files of type raw. **? TROUBLESHOOTING**

**Table 2:** Imaging and FCS settings to be used for green and red fluorescent proteins and dyes. The settings are given for Zeiss LSM microscopes.

|  |  |
| --- | --- |
| **Alexa-488/GFP settings** |  |
| **FCS APD detectors** |  |
| Main beam splitter (MBS) | 488, 488/561, 488/561/633 |
| Second beam splitter (NFT) | NFT 565 |
| Green channel emission filter (channel 2) | 505-540, 505-550 |
| **FCS GaAsP detectors** |  |
| Main dichroic | 488, 488/561, 488/561/633 |
| ChS1 | 501-544 |
| **Zeiss LSM, Imaging detectors** |  |
| Main dichroic | 488, 488/561, 488/561/633 |
| Ch1/ChS1 | 492-544 |
| **Alexa-568/mCherry settings** |  |
| **FCS APD detectors** |  |
| Main beam splitter (MBS) | 488/561, 488/561/633 |
| Second beam splitter (NFT) | NFT 565 |
| Red channel emission filter (channel 1) | 600-650 |
| **FCS GaAsP detectors** |  |
| Main beam splitter (MBS) | 488/561, 488/561/633 |
| ChS2 | 595-655 |
| **Zeiss LSM, Imaging detectors** |  |
| Main dichroic | 488, 488/561, 488/561/633 |
| Ch2/ChS2 | 595-655 |

**Table 3:**
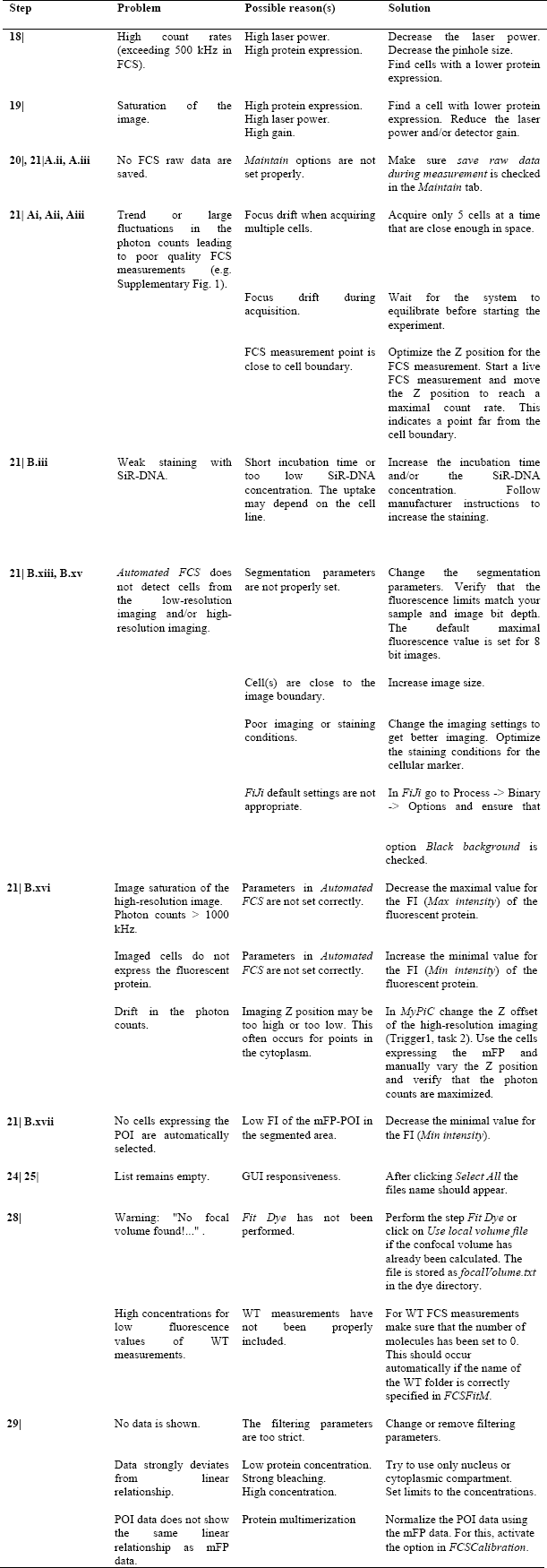
Troubleshooting table.

| Step | Problem | Possible reason(s) | Solution |
| --- | --- | --- | --- |
| 18\| | High count rates (exceeding 500 kHz in FCS). | High laser power.<br>High protein expression. | Decrease the laser power.<br>Decrease the pinhole size.<br>Find cells with a lower protein expression. |
| 19\| | Saturation of the image. | High protein expression.<br>High laser power.<br>High gain. | Find a cell with lower protein expression. Reduce the laser power and/or detector gain. |
| 20\|, 21\|A.ii, A.iii | No FCS raw data are saved. | *Maintain* options are not set properly. | Make sure *save raw data during measurement* is checked in the *Maintain* tab. |
| 21\| Ai, Aii, Aiii | Trend or large fluctuations in the photon counts leading to poor quality FCS measurements (e.g. Supplementary Fig. 1). | Focus drift when acquiring multiple cells. | Acquire only 5 cells at a time that are close enough in space. |
|  |  | Focus drift during acquisition. | Wait for the system to equilibrate before starting the experiment. |
|  |  | FCS measurement point is close to cell boundary. | Optimize the Z position for the FCS measurement. Start a live FCS measurement and move the Z position to reach a maximal count rate. This indicates a point far from the cell boundary. |
| 21\| B.iii | Weak staining with SiR-DNA. | Short incubation time or too low SiR-DNA concentration. The uptake may depend on the cell line. | Increase the incubation time and/or the SiR-DNA concentration. Follow manufacturer instructions to increase the staining. |
| 21\| B.xiii, B.xv | *Automated FCS* does not detect cells from the low-resolution imaging and/or high-resolution imaging. | Segmentation parameters are not properly set. | Change the segmentation parameters. Verify that the fluorescence limits match your sample and image bit depth. The default maximal fluorescence value is set for 8 bit images. |
|  |  | Cell(s) are close to the image boundary. | Increase image size. |
|  |  | Poor imaging or staining conditions. | Change the imaging settings to get better imaging. Optimize the staining conditions for the cellular marker. |
|  |  | *FiJi* default settings are not appropriate. | In *FiJi* go to Process -> Binary -> Options and ensure that option *Black background* is checked. |
| 21\| B.xvi | Image saturation of the high-resolution image. Photon counts > 1000 kHz. | Parameters in *Automated FCS* are not set correctly. | Decrease the maximal value for the FI (*Max intensity*) of the fluorescent protein. |
|  | Imaged cells do not express the fluorescent protein. | Parameters in *Automated FCS* are not set correctly. | Increase the minimal value for the FI (*Min intensity*) of the fluorescent protein. |
|  | Drift in the photon counts. | Imaging Z position may be too high or too low. This often occurs for points in the cytoplasm. | In *MyPiC* change the Z offset of the high-resolution imaging (Trigger1, task 2). Use the cells expressing the mFP and manually vary the Z position and verify that the photon counts are maximized. |
| 21\| B.xvii | No cells expressing the POI are automatically selected. | Low FI of the mFP-POI in the segmented area. | Decrease the minimal value for the FI (*Min intensity*). |
| 24\| 25\| | List remains empty. | GUI responsiveness. | After clicking *Select All* the files name should appear. |
| 28\| | Warning: "No focal volume found!..." . | *Fit Dye* has not been performed. | Perform the step *Fit Dye* or click on *Use local volume file* if the confocal volume has already been calculated. The file is stored as *focalVolume.txt* in the dye directory. |
|  | High concentrations for low fluorescence values of WT measurements. | WT measurements have not been properly included. | For WT FCS measurements make sure that the number of molecules has been set to 0. This should occur automatically if the name of the WT folder is correctly specified in *FCSFitM*. |
| 29\| | No data is shown. | The filtering parameters are too strict. | Change or remove filtering parameters. |
|  | Data strongly deviates from linear relationship. | Low protein concentration.<br>Strong bleaching.<br>High concentration. | Try to use only nucleus or cytoplasmic compartment. Set limits to the concentrations. |
|  | POI data does not show the same linear relationship as mFP data. | Protein multimerization | Normalize the POI data using the mFP data. For this, activate the option in *FCSCalibration*. |

### FCS and imaging data acquisition for calibration

21| FCS and imaging data can be obtained either by manual acquisition (Option A) or by automated adaptive feedback acquisition (Option B). See the Experimental Design section for more details.

A **Manual acquisition TIMING 0.5h setup, 2h measurement**

i. *Background measurements (5-10 cells).* Start the VBA macro *FCSRunner*(**Supplementary Software 1**). Specify the output folder as the *Calibration/WT* folder. Define the number of measurement points per cell. By default each point is associated to the cellular compartment nucleus/chromatin or cytoplasm (consult the software documentation for further details). For each compartment at least one point should be measured per cell. Focus in the well with WT cells and search for cells. Set the FCS and imaging settings according to the settings determined in 18| and 19|, 30 sec one repetition. Start the live mode and focus in XYZ to the cell of interest; add points to *FCSRunner* for the different compartments. Add up to 5 cells. Press `Image and FCS` in the *FCSRunner* VBA macro to start the acquisition. The VBA macro automatically acquires and saves images, FCS measurements, and FCS measurement coordinates at each cell position. Wait for the acquisition to finish. Delete the positions in *FCSRunner* and add new positions if more cells are needed. **CRITICAL STEP** When adding FCS points to the same object/cell, do not move the XY stage or change the focus. **? TROUBLESHOOTING**
ii. *Measurement of the mFP for the FCS calibration curve (10-20 cells).* Remove previous stage positions from the *FCSRunner.* Specify the output folder *Calibration/mFP* in *FCSRunner*. Move the stage to the well containing the cells expressing the mFP. Search for cells exhibiting low mFP expression levels. Switch to the FCS and imaging parameters determined in 18| and 19|, respectively, 30 sec one repetition. Verify that the image does not saturate and that the total photon counts are low (< 1000 kHz). Start the live mode and focus in XYZ to the cell of interest; add measurement points to *FCSRunner* for the different compartments. Add up to 5 cells. Press `Image and FCS` in the *FCSRunner* VBA macro to start the acquisition. Wait for the acquisition to finish. Delete the positions in *FCSRunner* and search for more cells to measure. Measure up to 20 cells. Make sure that the FCS raw data are saved. **CRITICAL STEP** The APD and GaAsP detectors are sensitive detectors. The protein expression level needs to be low in order to avoid saturation of the imaging intensities and exceeding of the FCS counts beyond 1000 kHz. Make sure that the imaging settings correspond to the imaging settings for the protein of interest. When adding FCS points to a cell do not move the XY stage or the focus. **? TROUBLESHOOTING**
iii. *Measurement of the fluorescently tagged POI (10-20 cells).* Remove previous stage positions in *FCSRunner.* Specify the output folder *Calibration/POI* in the *FCSRunner.* Move the stage to the well containing the cells expressing the fluorescently tagged POI. Search for cells that express the fluorescently tagged POI. Switch to the FCS and imaging parameters determined in 18| and 19|, respectively, 30 sec one repetition. Verify that the image does not saturate and that the photon counts are low (< 1000 kHz). For proteins tagged at their endogenous locus the expression levels are typically homogeneous within the population. In this case the previous step can be omitted. Start the live mode and focus in XYZ to the cell of interest; add measurement points to *FCSRunner* for the different compartments. Add up to 5 cells. Press 'Image and FCS' in the *FCSRunner* VBA macro to start the acquisition. Wait for the acquisition to finish. Delete the positions in *FCSRunner* and search for more cells to measure. Measure up to 20 cells. Make sure that the FCS raw data are saved. **CRITICAL STEP** Avoid acquiring FCS data in positions where the protein is bound to immobile structures (e.g. nuclear membrane). Indication of a large stable fraction is a strong bleaching during the FCS data acquisition. **? TROUBLESHOOTING**
B **Automated adaptive feedback acquisition (optional) TIMING 1.5 h setup (first time only), 2 h unattended acquisition**

i. *Loading the MyPiC ZEN Macro.* Start the macro (**Supplementary software 2**). Click on *Saving* and specify the output directory to *Calibration* directory (see 16|). Click on the *JobSetter* button.
ii. *Starting the Automated FCS Fiji plugin.* Start *FiJi* and import the automated FCS Fiji plugin (**Supplementary software 3**). Run the plugin by navigating to *Plugins > EMBL > Automated FCS.*
iii. *High-resolution imaging settings. HR.* These imaging settings are used for computing the FCS calibration curve and set the FCS measurement points using image analysis. Add an imaging channel for the cellular marker to the settings from 19|. Click the + button on the *JobSetter* to add the imaging job to *MyPiC* and name the job HR (high-resolution). **? TROUBLESHOOTING**
iv. *Low-resolution imaging settings. LR.* These imaging settings are used to automatically detect cells to be imaged and measured with FCS. Change the zoom settings in B.iii to acquire a large field of view. Adjust the pixel size and dwell time to achieve fast acquisition. The quality of the image should be good enough to allow for segmentation of the cellular DNA marker by *Automated FCS.* Use a Z stack if this improves the segmentation quality of the cellular marker. Click the + button in the *JobSetter* to add the imaging job to *MyPiC* and name the job LR (low-resolution).
v. *Autofocus imaging settings. AF.* To speed up the acquisition and minimize the hardware load it is recommended to keep the same light path settings as in B.iv. Change the settings in B.iv to XZ line-scanning and reflection mode (see also the software manual for **Supplementary software 2**). Choose a laser line that is not reflected by the main beam splitter (MBS) but detected with the current detector settings. For example use the Argon 514 nm laser line with the MBS 488/561/633 and GFP detection range. Set the laser power and gain to achieve a visible reflection without saturating the detector. Set the number of stacks to cover 20-80 μm with a 100-500 nm Z step size. A small Z step size gives higher precision but requires a longer acquisition time in the absence of a piezo Z stage. Verify that the imaging yields a thin bright line. Click the + button in the *JobSetter* and name the job AF (Autofocus). **CRITICAL STEP** With the recommended MBS and laser settings the transmitted light can be high and damage the detector. The user should start with a low laser light and gain and then adjust it accordingly.
vi. *FCS settings.* Set in ZEN the FCS settings for the mFP-POI (see also 18|). Use one repetition and 30 sec measurement time. In the *JobSetter* click on the FCS tab. Load the FCS settings from ZEN by clicking on the + button and name the job POIFCS. ZEN will prompt the user to save the light path settings.
vii. *Task 1 of the Default pipeline: Autofocus.* Click on the *Default* button in *MyPiC.* Click the + button and add the AF-job as first task to the default pipeline with a double click. Set *Process Image/Tracking* of the first task to *Center of mass (thr)* and click on *TrackZ*. At each position and repetition *MyPiC* computes the fluorescence center of mass of the upper 20% of the signal and updates the Z position accordingly.
viii. *Task 2 of the Default pipeline: Low-resolution imaging.* Click the + button of *MyPiC* and add the LR job as second task to the default pipeline with a double click. Test the default pipeline by pressing the play button in the *Pipeline Default tasks* frame. The image is stored in the directory *Test* with the name DE_2_W0001_P0001_T0001.lsm|czi. Adjust the value of the Z offset for the LR-job to achieve the necessary imaging position from the cover glass. Acquire a final test image. Set *Process Image/Tracking* of the second task to *Online Image Analysis.* At every position and repetition *MyPiC* first acquires the autofocus image followed by a low-resolution image. Then *MyPiC* waits for a command from the *Automated FCS* FiJi plugin (B.xii).
ix. *Task 1 of the Trigger1 pipeline: Autofocus.* Press the *Trigger1* button in *MyPiC.* Press the + button and add the AF job as first task to the Trigger1 pipeline with a double click. Set *Process Image/Tracking* of the first task to *Center of mass (thr)* and click on *TrackZ*.
x. *Task 2 of the Trigger1 pipeline: High-resolution imaging.* Press the + button in *MyPiC* and add the HR job as second task to the Trigger1 pipeline with a double click. Test the Trigger1 pipeline by pressing the play button in the *Pipeline Trigger1 tasks* frame. The image is stored in the directory *Test* with the name TR1_2_W0001_P0001_T0001.lsm|czi. Adjust the value of the Z offset for the HR job to achieve the necessary imaging position from the cover glass. Acquire a final test image. Set *Process Image/Tracking* of the second task to *Online Image Analysis.*
xi. *Task 3 of the Trigger1 pipeline: FCS acquisition.* Press the + button of *MyPiC* and add the POIFCS job as third task to the Trigger1 pipeline.
xii. *Automatically detect the cell of interest in the Default pipeline.* In the *FiJi* macro *Automated FCS* click on *Parameter Setup.* In the *Job1* column change *Pipeline* to *Default.* Change *Task* to *2* and *Command* to *trigger1.* Specify the channel of the cellular marker in *Channel segmentation.* Set in *Number of particles* the maximum number of cells to process to 4. Specify in *Pick particle* how the cells should be chosen: use *random.* Please refer to the manual of *Automated FCS* (**Supplementary Software 3**) for a detailed explanation of the software options.
xiii. *Adjust image analysis settings to detect the cell of interest in the Default pipeline.* Test cell detection by clicking *Run on file.* If cells of interest are detected, an image with the processing results will be generated. To improve segmentation of the cellular marker and separate single cells, the user can change the thresholding method *(Seg. method)* and area size limits. In *Channel intensity filter1* specify the channel of the cellular marker and set a lower and upper value for the mean FI. This avoids cells with an abnormally high FI of the cellular marker (e.g. apoptotic cells). Test the settings with the *Run on file* command. In *Channel intensity filter2* specify the channel of the FP and set a lower and upper value for the mean FI. The lower value should be set so that cells not expressing the FP are avoided. The upper value avoids saturation in the high-resolution image and FCS measurements. Test cell detection by clicking *Run on file* and select the image generated in B.viii. Acquire additional images of cells expressing the mFP and endogenously mFP-POI to further test the image analysis settings. **? TROUBLESHOOTING**
xiv. *Automatically place FCS measurements in the Trigger1 Pipeline.* In the *Job2* column of the *Parameter Setup* window change *Pipeline* to *Trigger1.* Change *Task* to *2* and *Command* to *setFcsPos.* Specify the cellular marker channel in *Channel segmentation.* Specify the number of FCS measurement points and the pixel distance from the boundary of the segmented object *(FCS pts. region1/2* and # *oper. (erode <0, dilate >0),* respectively). Use at least one point inside (*# oper* < 0, nucleus) and one point outside (*# oper* > 0, cytoplasm) of the segmented cell marker.
xv. *Adjust image analysis settings to place FCS points in the Trigger1 pipeline*. Verify detection of cells and the placement of FCS points by clicking *Run on file* in *Parameter Setup* and selecting the image generated in B.x. If objects of interest are detected, an image of the processing results will be generated. As for step B.xiii modify the *Seg. method,* the area size limits, the *Channel intensity filter1*and *Channel intensity filter2* if required. If necessary, adjust the pixel distance to the object border to ensure correct placement of the FCS measurement points *(# oper. (erode <0, dilate >0)).* Acquire additional images of cells expressing the mFP and mFP-POI to further test the image analysis settings. **? TROUBLESHOOTING**
xvi. *Automated measurement of the FP (10-20 cells).* In *MyPiC* specify the output directory to *Calibration/mFP*. Press *Default Positions,* click on *Multiple* and mark 3-10 positions in the mFP well. To improve the throughput manually select positions where several cells in the field of view express the mFP construct. In *Automated FCS* set the *Directory to monitor* to *Calibration/mFP*. Press start in *Automated FCS* and *MyPiC.* When the acquisition is finished press *Stop* in *Automated FCS.* **CRITICAL STEP** When this step is performed for the first time the user should supervise the acquisition to ensure that the cells obtained from the analysis express the FP at the appropriate level. Furthermore, images should not be saturated and the FCS counts should be below 1000 kHz. See also troubleshooting on how to adjust the settings in *Automated FCS* (step B.xv). **? TROUBLESHOOTING**
xvii. *Automated measurement of the fluorescently tagged POI (10-20 cells).* In *MyPiC* specify the output directory to *Calibration/POI.* Press *Default Positions,* delete previous positions and mark 3-10 positions in the POI well. To improve the throughput, manually select positions where several cells in the field of view express the mFP-POI. In *Automated FCS* specify the current directory to *Calibration/POI.* Press start in *Automated FCS* and *MyPiC.* When the acquisition is finished press *Stop* in *Automated FCS.* **CRITICAL STEP** Same considerations as for step B.xvi apply for this step. **? TROUBLESHOOTING**
xviii. *Automated background measurements* (5-10 cells). In *MyPiC* specify the output directory to *Calibration/WT.* Press *Default Positions,* delete previous positions and mark 2-3 positions in the WT well. In *Automated FCS* set the current directory to *Calibration/WT.* Change *Channel intensity filter2* to *None.* Press start in *Automated FCS* and start in *MyPiC.* When the acquisition is finished press *Stop* in *Automated FCS.*

22| *(Optional) Additional imaging.* Perform additional imaging using the same detector and laser settings as specified in 19|. The FI in these images can be converted to concentrations and protein number using the calibration parameters (see step 30|). **CRITICAL STEP** To convert fluorescence intensities to protein numbers the image needs to be a 3D (XYZ) or 4D (XYZT) stack.

**Data Processing TIMING 10 min setup, 0.5-1h of unattended computation (depending on the number of measurements), 30 min quality control, 0.5-1h image processing**

23| Start the tool FCSFitM (**Supplementary Software 4**). **CRITICAL STEP** See the manual of the software for detailed explanations.
24| *Computing the background photon counts with FCSFitM.* Change to tab *FCS background* in *FCSFitM.* Specify the directory *Calibration/WT.* Press *SelectAll* and *Compute* to calculate the average photon counts for all measurements. For the background value no distinction is made between measurements in the nucleus and the cytoplasm. Write down the counts of Ch1 (or Ch2). This will be needed in step 26|. **CRITICAL STEP** In this and all subsequent analysis steps Ch1 indicates the channel with the lowest wave length (e.g. GFP, Alexa488) and Ch2 the channel with the higher wave length (e.g. mCherry, Alexa568). If APDs are used the ZEN notation of Ch2 and Ch1 corresponds in the analysis to Ch1 and Ch2, respectively. **? TROUBLESHOOTING**
25| *Effective confocal volume estimation from the fluorescent dye using FCSFitM.* Change to tab *Fit Dye.* Specify the diffusion coefficient of the fluorescent dye used for Ch1 or Ch2. The default values are for Alexa488 and Alexa568 at 37 °C. See also **Supplementary Note 2**. Specify the directory that contains the fluorescent dye data *(Calibration/dye).* Press *SelectAll* and *Compute.* Wait until the computation is finished. The program computes a best estimate for *w_0_* and *κ* by performing repeated rounds of parameter optimization. Finally a mean effective confocal volume *V_eff_* is computed (**Supplementary Note 1** and **Box 2**). Check that results have been saved in *Calibration/dye/focalVolume.txt* and *Calibration/dye/optimisedFit.* **? TROUBLESHOOTING**
26| *Computing correlations and corrections using FA.* Start FA. Specify the raw data format *Zeiss/ZEN (*.zen).* Specify the import settings according to the used detectors. See also the manual for the **Supplementary Software 4**. Select the path to *Calibration* and click *Include subdirectories.* Add all measurements and name the session *2c* for later reuse. Press *Check files* and wait for all files being processed. In *Modify and correlate* set the base frequency to 100.000 Hz (i.e. 10 μs time interval). Press *Calculate all* and wait for the end of the computation. In the *Intensity corrections* tab enter the background values obtained from 24| in the *Offset Ch1* text element. Press *Apply to* and *Calculate All.* Wait for the computation to finish. Perform a visual quality check (QC) of the data according to the requirements in Supplementary Fig. 1 and annotate traces that do not pass the QC with a ‘x’. Parameter-based QC are applied at step 29|. In the *Save, export and report* tab press *Save all* button and the *FA format* button. This saves the data to the tab delimited result table *Calibration/2c.res* (see also documentation to **Supplementary Software 4**). **CRITICAL STEP** On a Zeiss Confocor, to maintain the channel convention (see 24|) when loading data acquired using APDs you may need to swap the order of the channels (see also documentation to **Supplementary Software 4**).
27| *Extracting fluorescence intensities at the FCS measurement points.* Start *FiJi.* Start the macro *Plugins* > *EMBL* > *FCSImageBrowser*. Press *Load.res file* and load the result table created in 26|. Click on *Report* to generate the table *2c.fint.* The table contains the fluorescence intensities in ROIs of different sizes at the FCS measurement points. If images have been acquired using the adaptive feedback pipeline, perform an additional visual QC of the images. Omit points located at the cell boundary. Annotate the FCS points that do not pass the visual QC of the images with a ‘x’ in *FA.* In the *Save, export and report* tab press *Save all.* Press *FA format* button to save the updated results table.
28| *Fitting the ACFs functions using FCSFitM.* Press *Add res file* and load the result table from 26| into *FCSFitM.* Press *Run fit* to perform fits to the autocorrelation data. The program generates two separate tables, *1c opt.res* and *2c_opt.res* for one- and two-component anomalous diffusion models, respectively. *FCSFitM* uses the data from 25| and 27|. The generated tables summarize the fit parameters, the concentrations, and fluorescence intensities at the FCS measurement points (see documentation of **Supplementary Software 4**). The ACF fit results can be viewed in *FA* using the session name *1c_opt* or *2c_opt (1c_opt_w* and *2c_opt_w* if a weighted fit is used). **? TROUBLESHOOTING**
29| *Computing FCS calibration curve using FCSCalibration.* All data to compute the calibration curve are stored in the tables generated in step 28|. The *R* application *FCSCalibration* provides a graphical interface to further QC the data and automatically computes the calibration curve. Start the *FCSCalibration* and specify the directory containing the *2c_opt.res* (step 28|). If required, change the name of Dye, *POI,* and *mFP* entries to match the name of the directories of the current experiment. Choose the ROI size and the compartment (nucleus, cytoplasm or all) to use. Choose the filtering parameters by investigating their distribution. Typically the listed default parameters work for most applications. Press *Report* to save the calibration data to the folder *Results/Calibration*. **? TROUBLESHOOTING**
30| *Converting fluorescence intensities to concentrations and protein numbers.* For the linear transformation of the fluorescence intensities to concentration the user only needs the parameters of the FCS calibration curve step 29|. To convert concentrations into number of proteins the voxel volume is required, which is stored in the image metadata. Conversion in protein number per pixel is only performed for 3D and 4D data. The FiJi plugin *FCSCalibrateImage* (**Supplementary Software 4**) reads the image, the metadata, and the calibration data to generate a calibrated image for the channel of interest.

## TIMING

Steps 1|-9|: Sample preparation: 1 h hands-on, 1 d waiting time, 15 min hands-on.

Steps 10|-20|: Microscope setup and effective confocal volume measurement: 1 h to 2 h depending on the experience.

Step 21| Option A: Manual acquisition: 0.5 h setup, 2 h measurement.

Step 21| Option B: Automated adaptive feedback acquisition: 1.5 h setup (first time only) and 2 h unattended acquisitions. When settings are reused, the setup time reduces to less than 10 min.

Step 22|: The time required for acquiring additional imaging depends on the application (2D, 3D or 4D imaging).

Steps 23|-30|: Data Processing: 10 min setup, 0.5-1 h of unattended computation (depending on the number of measurements), 30 min quality control, 0.5-1 h image processing.

Box 1: Automated FCS-calibrated imaging: 2-3.5 h.

Box 2: Estimating the effective confocal volume using a fluorescent dye: Acquisition and data processing 30 min.

Box 3: Generating dextran labeled with a fluorescent dye: Preparation 3 h and overnight dialysis.

## ANTICIPATED RESULTS

The workflow described in this protocol provides a calibration coefficient for converting FI data to protein concentrations and numbers. The coefficient needs to be calculated for every experiment. Depending on the stability of the system, its value may remain similar over time. In combination with further image analysis steps using image analysis software such as MATLAB or ImageJ the stoichiometry of the protein of interest on cellular structures can be computed (Fig. 3). The adaptive feedback imaging workflow can be used to acquire additional FCS measurements and 4D image movies in high-throughput. The FCS measurements can be further used to study the biophysical properties of the POI (diffusion coefficient, aggregation). In our laboratory we used this method to generate FCS-calibrated 4D image movies of cells during mitosis ^12,15^.

Figure 3 shows an example for the quantification of the protein density on the nuclear envelope. A cell line expressing NUP107, a nucleoporin which is a constitutive part of the NPC, endogenously tagged with mEGFP has been used. Based on a DNA marker the nucleus was segmented in 3D to extract the localization of the nuclear envelope. For further processing the equatorial plane of the nucleus has been used (Fig. 3b) and the mean FI in the nuclear rim has been extracted. Depending on imaging settings and microscope, the value of the arbitrary fluorescent intensity units on the nuclear envelope can be significantly different (Fig. 3c). However, after conversion to absolute protein numbers using a separate calibration curve for each microscope, comparable results between the different imaging systems are obtained (Fig. 3d). The quantification of NUP107 using this method yielded a protein density at the nuclear envelope that is in good agreement with the density predicted from the NUP107 stoichiometry (32 proteins/NPC, ^24^) and the NPC density in interphase (10.66 ± 2.69 NPCs/μm^2^, ^57^) (Fig. 3d, gray shaded area).

## ACKNOWLEDGMENTS

We thank the mechanical and the electronics workshop of EMBL for custom hardware, the Advanced Light Microscopy Facility of EMBL for microscopy support and the Flow Cytometry Core Facility of EMBL for cell sorting. We gratefully acknowledge Bianca Nijmeijer, Stephanie Alexander and Arina Rybina for critically reading the manuscript. We thank Thomas Ohrt (Zeiss) for support in the hardware control. This work was supported by grants to J.E. from the European Commission EU-FP7-Systems Microscopy NoE (Grant Agreement 258068), EU-FP7-MitoSys (Grant Agreement 241548) and iNEXT (Grant Agreement 653706), as well as by EMBL (A.Z.P., Y.C., N.W., M.J.H., B.K., M.W., J.E.). Y.C. and N.W. were supported by the EMBL International PhD Programme (EIPP).

## AUTHOR CONTRIBUTIONS

A.Z.P. performed the experiments, the analysis and designed the software packages. A.Z.P with the help of Y.C. and M.W. developed the protocol. N.W. tested the protocol and helped in writing the software manuals. B.K. created the homozygous cell line and provided the dextran labelling protocol. M.J.H. provided the code to segment the cells. A.Z.P. with the help of N.W., B.K., and J.E. wrote the protocol.

## Box 1: Automated FCS-calibrated imaging.

Timing 2-3.5 h. This box gives an outline of the requirements for acquiring FCS and imaging data automatically. The two supplementary softwares *MyPiC* (**Supplementary Software 2**) and *Automated FCS* (**Supplementary Software 3**) are used. With an adaptive feedback workflow ^17,18^ cells are automatically selected according to their morphology and protein expression level, imaged and measured using FCS. This yields reproducible measurements and a higher throughput. The procedure requires a fluorescent marker, such as a DNA marker, or any other segmentable fluorescent marker, to identify cells. If the signal of the POI is sufficiently good, no additional markers are required. To perform automated acquisition three different imaging settings are used:

- A high-resolution imaging of the fluorescent marker and FP to place FCS measurements and acquire the reference image (Step 21| B.iii).
- A low-resolution imaging of the fluorescent marker and FP to identify cells of interest (Step 21| B.iv).
- An autofocus XZ scan to detect the cover glass reflection and correct for drift in Z (Step 21| B.v).

Refer to the corresponding manuals for **Supplementary Software 2-3** for a detailed description. After the first setup settings can be reused. This reduces the setup time to less than 10 min.

## Box 2: Estimating the effective confocal volume using a fluorescent dye.

Timing 30 min. This box describes how to compute the parameters for determining the effective confocal volume. For FCS-calibrated imaging this step is particularly important since it is used to scale all measurements. Use a fluorescent dye in aqueous solution (5-50 nM, ^60^) with high molecular brightness, spectral match with available laser lines and filters, spectral match with the FP of interest, and a known diffusion coefficient. To investigate proteins tagged with green/yellow FPs (e.g. GFP, YFP, mNeonGreen), Alexa488 is a suitable dye. For proteins tagged with a red FP (e.g. mCherry, mScarlet), Alexa568 is suited.

Procedure:

1. Adjust the high NA water immersion objective to the thickness of the cover slip glass by turning the correction ring (Fig. 2a).
2. Assess the optimal adjustment by measuring the counts per molecule (CPM), the photon counts, and the reflection at the immersion water glass interface in a XZ scan. At the optimal adjustment, CPM and photon counts are maximal and the reflection line is the thinnest (Fig. 2a).
3. Acquire repeated FCS measurements (up to 6, 30 sec) of the reference dye.
4. Fit the ACFs (Fig. 2b) to a one component model of diffusion (see also **Supplementary Notes 1** and **2**)

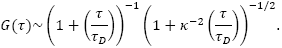
5. Obtain the diffusion time *τ_D_* from the fit of the ACFs, that is the average time the fluorophore remains in the confocal volume, and the ratio of axial to lateral radius of the confocal volume *к* (typically between 4-8). Use the diffusion time to compute the lateral radius

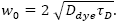 The diffusion coefficients *D_dye_* for Alexa488 and Alexa568 at 27 °C are *D_A488_* = 3 6 5 ± 20 μm^2^/sec and *D_A568_* = 410 ± 40 μm^2^/sec (M. Wachsmuth personal communication and ^61^) yielding at 37 °C a mean of *D_A488_* = 464.23 μm^2^/sec and *D_A568_* = 521.46 μm^2^/sec (**Supplementary Note 2**, Eq. S7). Diffusion coefficients for additional fluorescent dyes can be found in ^62^. Typical values for *w_0_* are 190-250 nm.
6. Determine the effective confocal volume by

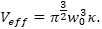

The processing and fitting steps 4-6 can be performed using *FCSFitM* (**Supplementary Software 4**).

## Box 3: Generating dextran labeled with a fluorescent dye.

Timing 3 h and overnight dialysis.

This box describes the protocol to label dextran with a fluorescent dye. The reagent can be used to stain the extracellular medium and thereby provides a marker for segmenting the cellular boundary.

Procedure:

1. Dissolve 125 mg dextran, Amino, 500 000 MW, in 6.25 ml of 0.2 molar NaHCO_3_ (pH 8.0-8.3, freshly prepared).
2. Mix well by stirring.
3. Dissolve 10 mg Dy481XL-NHS-Ester (Dyomics 481XL-01) in 0.7 ml of DMSO.
4. Add the Dy481XL solution directly into the dextran solution (from step 1) as quickly as possible since it is not stable.
5. Mix at RT, stirring with ∼500 rpm for 1 h.
6. Use 3x PD-10 Sephadex desalting columns to separate unreacted Dy481XL following the suppliers manual and elute with 3.5 ml water per column.
7. Dialyze against water (sterile MonoQ) using Slide A-Lyzer 10000 MWCO at 4 °C. Exchange the water after ∼2 h. Further dialyze overnight.
8. Concentrate the labeled dextran using Viva Spin 30.000 PES as described in the supplier’s manual.
9. Measure the amount at the wavelength of 515 nm and calculate the concentration of the fluorescently labeled dextran using the following equation:

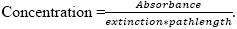
10. Aliquot and store at - 20 °C. The reagent can be stored for > 1 year.

## Supplementary material

**Supplementary Software 1: Microscopy software to manually acquire FCS-calibration data.**

**Supplementary Software 2: Microscopy software to automatically acquire FCS-calibration data.**

**Supplementary Software 3: Analysis software for automatic acquisition of FCS-calibration data.**

**Supplementary Software 4: Analysis software to process the FCS and imaging data. Supplementary Figure 1: Quality control of FCS traces.**

**Supplementary Figure 2: CPM and effective volume dependency on pinhole diameter and laser power.**

**Supplementary Figure 3: Simulated FCS-calibrated imaging.**

## Combined Supplementary Information file

**Supplementary Table 1: Software packages for FCS-calibrated imaging. Supplementary Table 2: Parameters of the diffusion models.**

**Supplementary Note 1: Effective confocal volume.**

**Supplementary Note 2: Fitting of the ACFs.**

**Supplementary Note 3: Correction for background and photobleaching. Supplementary Note 4: Estimate the number of proteins on small structures. Supplementary Note 5: Total fluorophores in a point source.**

**Supplementary Note 6: Density of fluorophores on a membrane.**

## Supplementary material legends

### Supplementary Software 1: Microscopy software to manually acquire FCS-calibration data

Contains the VBA macro *FCSRunner.lvb* for acquiring imaging and FCS data to later compute the FCS calibration parameters. The macro runs with ZEN black edition (version ≥ 2010) on Zeiss LSM microscopes. Please read the documentation located in the *wiki* directory or the Wiki at https://git.embl.de/grp-ellenberg/fcsrunner/wikis for detailed information on how to operate the software. In order to keep track of possible software updates, the user can clone the package from the primary public software repository using the following *git* command: *git clone https://git.embl.de/grp-ellenberg/fcsrunner.git.*

### Supplementary Software 2: Microscopy software to automatically acquire FCS-calibration data

Contains the VBA macro *MyPiC.lvb.* The macro runs with ZEN black edition (version ≥ 2010) on Zeiss LSM microscopes. *MyPiC* is used to perform complex imaging and FCS workflows with the option to integrate online image analysis (see **Supplementary software 3**). The latter feature can be used to automatically acquire a large set of imaging and FCS data for the FCS calibration curve. Please read the documentation located in the *wiki* directory or the Wiki at https://git.embl.de/grp-ellenberg/mypic/wikis for detailed information on how to install and operate the software. In order to keep track of possible software updates, the user can clone the package from the primary public software repository using the following *git* command: *git clone https://git.embl.de/grp-ellenberg/mypic.git.*

### Supplementary Software 3: Analysis software for automatic acquisition of FCS-calibration data

FiJi plugin *AutomatedFCS.py* and related software. The plugin allows performing online image analysis for images acquired using *MyPiC.* The plugin performs intensity based segmentation in combination with a watershed step to separate merging objects. The segmented objects are then used to define new coordinates for imaging and/or fluorescence correlation measurements. Please read the documentation located in the *wiki* directory or the documentation at https://git.embl.de/grp-ellenberg/adaptive_feedback_mic_fiii/wikis for detailed information on how to install and operate the software. In order to keep track of possible software updates, the user can clone the package from the primary public software repository using the following git command: *git clone https://git.embl.de/grp-ellenberg/adaptive_feedback_mic_fiji.git.*

### Supplementary Software 4: Analysis software to process the FCS and imaging data

Analysis workflow for FCS-calibrated imaging. The software package contains*MATLAB-, R-*, and *FiJi*-based tools to extract fluorescence intensities, compute FCS calibration parameters and convert the fluorescence intensities to physical quantities. The package includes:

- *FCSImageBrowser* (in folder *fiji*) is a *FiJi* script to browse through the acquired images and extract the fluorescence intensities at the FCS measurement points.
- *FCSFitM* (in folder *matlab/FCSFitM*) is a *MATLAB* tool to batch fit the correlation data, compute the effective volume and the concentrations. For convenience a compiled Windows installer is included *(FCSFitM_web.exe).* The compiled version does not require a *MATLAB* license.
- *FCSCalibration* (in folder *R/FCSCalibration*) is a *shiny R* application to interactively perform quality control of the data and compute FCS calibration parameters. For convenience we include a packed version for Windows.
- *FCSCalibrateImage* (in folder *fiji*) is a *FiJi* script to convert fluorescence intensities to concentration and protein numbers.

Please read the documentation located in the *wiki* directory or the Wiki at https://sit.embl.de/gry-ellenbers/fcsanalyze/wikis for detailed information on how to install and operate the different elements of the software package. In order to keep track of possible software updates, the user can clone the package (without the compiled versions) from the primary public software repository using the following *git* command: *git clone https://git.embl.de/grp-ellenberg/fcsAnalyze.git.*

---

**Supplementary Figure 1:**
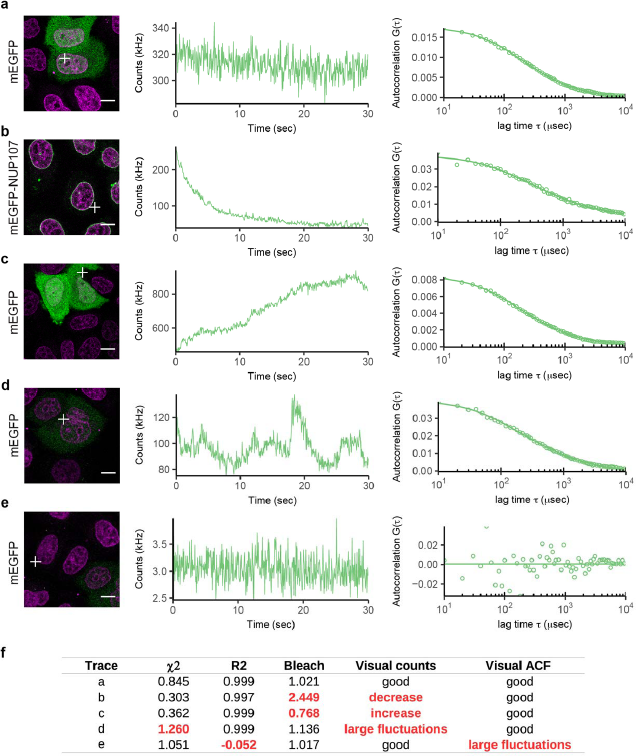
Quality control of FCS traces. (a) Typical trace that passes the quality control (QC) according to the parameters shown in f. The QC is based on thresholds applied to fitting parameters such as the sum of squared residuals *χ^2^*, the coefficient of variation *R^2^*, and properties of the photon counts traces (e.g. Bleach coefficient, as calculated by FA). A visual inspection of the photon counts and autocorrelation traces can also be used for quality control. Scale bar 10 μm. (b) FCS measurement point is at the boundary of the cytoplasmic compartment causing a decreasing drift in photon counts. (c) FCS measurement point is at the boundary of the cytoplasmic compartment causing an increasing drift in photon counts. (d) FCS measurement point is at the boundary between a dim and a brighter cell causing large fluctuations in the photon counts. (e) FCS measurement point is in a cell that does not express a fluorescent protein. (f) Table of parameters used for the QC. The measurements in b and c do not pass the QC according to the bleach parameter. The measurement in d does not pass the QC according to the *χ^2^* value. The measurement in e does not pass the QC according to the *R^2^* value. None of the traces in b-e pass the visual inspection. The thresholds used were *χ^2^* < 1.2, *R^2^* > 0.9, 0.8 < Bleach < 1.2. The thresholds can be interactively set in the *FCSCalibration* software (**Supplementary Software 4**).

**Supplementary Figure 2:**
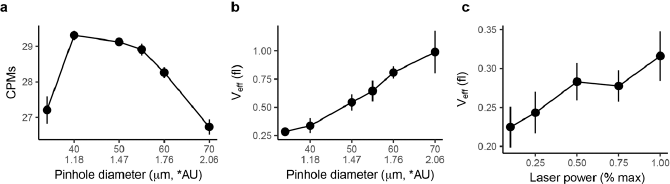
CPM and effective volume dependency on pinhole diameter and laser power. (a) Counts per molecule (CPM) as function of the pinhole diameter for Alexa488 in water. (b) Effective confocal focal volume estimated as described in the supplement (**Supplementary Notes 1-2** and **Box 2**) as function of the pinhole diameter for Alexa488. (c) Effective confocal focal volume as function of the excitation laser (Argon 488 nm). The pinhole was 34 μm.

**Supplementary Figure 3:**
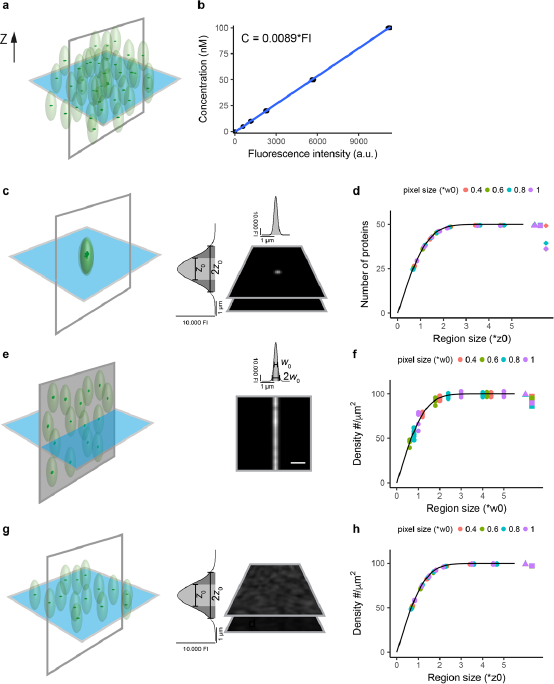
Simulated FCS-calibrated imaging. (a) Illustration of a homogeneous fluorophore distribution. (b) The slope of the fluorophore concentration as function of pixel FI gives the calibration coefficient. Simulated PSF is a 3D Gaussian (Eq. S1). The size of the PSF is characterized by its *e^2^* decay *w_0_* and *z_0_* in XY and Z direction, respectively. Here and in all subsequent panels background and detector noise are not simulated. (c) Simulated Z-stack of a point source (50 fluorophores). (d) Using the calibration coefficient and Eqs. S15, S16, and S19 the total number of proteins in each plane is calculated. Several planes along the Z-direction are summed (circles). The expected protein number is obtained when a large region along Z is considered (*>2*z0*). Simulated images have a pixel-size of 0.4, 0.6, 0.8, 1 times *w_0_* in all directions (different colors). Triangles give the result when all pixels are considered (Eq. S19). Squares give the approximation using the integral of the PSF along Z (Eq. S21). Diamonds give the approximation when the integral of the whole PSF and the pixel with the highest intensity is used (Eq. S23). (e) Simulated fluorophores distributed in a XZ plane (density of 100 fluorophores/μm^2^). Scale bar 10 μm.(f) Protein density is computed using Eqs. S15 and S24. The expected density is obtained for a region width along the X direction of > *2*w_0_*. Triangles give the result when all pixels are considered. Squares give the approximation when the integral of the PSF along X is used (Eq. S26). (g) Simulated fluorophores distributed in a XY plane (density of 100 fluorophores/μm^2^). (h) The average protein density is summed for planes along Z (Eqs. S15 and S17). The expected density is obtained for a region width > *2*z_0_*. Triangles give the result when all pixels are considered. Squares give the approximation when the integral of the PSF along Z is used (Eq. S29). Each fluorophore has a simulated intensity of 1000 (a.u.), the simulated PSF is characterized by *w*_0_ = 250 nm, *z*_0_ = 1500 nm. The black lines give the theoretically expected result after integration of the Gaussian function 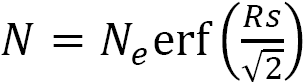 where *N_e_* is the expected protein number/density (50 in d, and 100 in f and h) and *Rs* is the region size in unit of the PSF characteristic size.

**Supplementary Table 1:** Software packages for FCS-calibrated imaging. Bundled packages are found on the Nature Protocols website. The links to the *git* repositories hosting the most recent source code is given. Recent bundled packages for **Supplementary Software 1-3** are found at https://git.embl.de/2rp-ellenberg/ followed by fcsrunner/tags,mypic/tags, and *adaptive_feedback_mic_fiii/tags,* respectively. Recent bundled versions of Fluctuation Analyzer 4G^18^ and **Supplementary Software 4** can be downloaded from *https://www-ellenberg.embl.de/resources/data-analysis.*

| Software tool | Software requirements | Operating system | Description | Source |
| --- | --- | --- | --- | --- |
| **FCSRunner**<br>Supplementary Software 1 | ZEN black edition ≥ 2010 | Windows | VBA ZEN macro to perform combined FCS and image acquisition. Install on data acquisition computer. | https://git.embl.de/grp-ellenberg/fcsrunner |
| **MyPiC**<br>Supplementary Software 2 | ZEN black edition ≥ 2010 | Windows | VBA ZEN macro to perform complex imaging and FCS pipelines. Automated data acquisition in combination with AutomtedFCS plugin. Install on data acquisition computer. | https://git.embl.de/grp-ellenberg/mypic |
| **Automated FCS**<br>Supplementary Software 3 | Fiji | Windows<br>(Linux, Mac OSX) | A Fiji plugin that interacts with MyPiC and allows among other things for unsupervised FCS data acquisition. Install on data acquisition computer. | https://git.embl.de/grp-ellenberg/adaptive_feed_back_mic_fiji<br>directory: *automatedfcs* |
| **FCSFitM**<br>Supplementary Software 4 | Matlab version ≥ R2014a<br>Toolboxes: Optimization, Statistics<br>Fluctuation Analyzer 4G | Windows, Linux,<br>Mac OSX | Matlab workflow to batch fit FCS data, compute effective focal volumes, and concentrations. As input data the software uses the correlation curves generated by Fluctuation Analyzer 4G. | https://git.embl.de/grp-ellenberg/fcsanalyze<br>directory: *matlab/FCSFitM* |
| **setup_FCSFitM_web.exe**<br>Supplementary Software 4 | Fluctuation Analyzer 4G | Windows 64-bit | The compiled version of FCSFitM for Windows. |  |
| **FCSImageBrowser**<br>Supplementary Software 4 | Fiji | Windows, Linux,<br>Mac OSX | Fiji plugin to visualize images and FCS positions and extract fluorescence intensities. | https://git.embl.de/grp-ellenberg/fcsanalyze<br>directory: *fiji* |
| **FCSCalibration**<br>Supplementary Software 4 | R (≥3.3.2)<br>RStudio (recommended)<br>the installation of package *fcsresfun* will force the installtion of missing packages | | R program to interactively compute FCS calibration parameters | https://git.embl.de/grp-ellenberg/fcsanalyze<br>directory: *R/FCSCalibration*<br>*R/fcsresfun* |
| **setup_FCSCalibration.exe**<br>Supplementary Software 4 | Fluctuation Analyzer 4G | Windows 64- bit | Bundled version of FCSCalibration. |  |
| **FCSCalibrateImage**<br>Supplementary Software 4 | Fiji | Windows, Linux,<br>Mac OSX | Fiji plugin to convert fluorescence intensities to concentrations and protein numbers using the FCS calibration parameters | https://git.embl.de/grp-ellenberg/fcsanalyze<br>directory: *fiji* |

**Supplementary Table 2:** Parameters of the diffusion models. Parameters of the models to fit the ACF of the protein and fluorescent dye (Eqs. S4-S5).

| Parameter name | Meaning |
| --- | --- |
| $N$ | Number of molecules in confocal volume |
| $\tau_{D_i}$ | Diffusion time of $i$-th component |
| $\alpha_i$ | Anomalous diffusion coefficient of $i$-th component |
| $f_1, f_2 = 1 - f_1$ | Fraction of each component |
| $\theta_T$ | Fraction of fluorophore in a non-fluorescent state |
| $\tau_T$ | Relaxation time of the non-fluorescent state |
| $\kappa = z_0/w_0$ | Structural parameter |

## Supplementary Note 1: Effective confocal volume

To compute the effective confocal volume we approximate the PSF by a 3D Gaussian

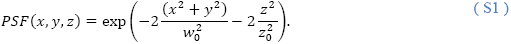

The parameters *w_0_* and *z_0_* characterize the *e^2^* decay length of the PSF. We also define the structural parameter as the ratio *κ = z_0_/w_0_.* The effective confocal volume for FCS^2^ is given by

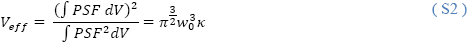

This is larger than the confocal volume given by

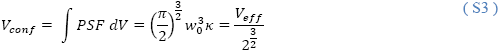

## Supplementary Note 2: Fitting of the ACFs

For fitting of the ACFs of the fluorescent proteins we assume a 3D Gaussian focal volume (Eq. (S1)) and use a two component anomalous diffusion model with fluorescent protein-like blinking ^1^

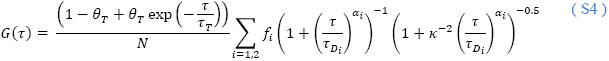

In the software *FCSFitM* (**Supplementary software 4**) the data is fitted using the MATLAB routine *lsqnonlin.* The meaning of the parameters is listed in Table S2. As expected for the number of proteins *N* we obtain only minor differences (less than 2%) between a one component (*f*_1_ = 1) or a two component model.

For the fluorescent dye in solution we use a single component non-anomalous diffusion model with triplet-like blinking

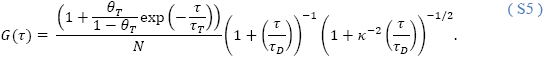

Equation (S5) is fitted to the ACF curves for the fluorescent dye to obtain the diffusion time *τ_D_* and the structural parameter *κ.* The width of the focal volume is then given by

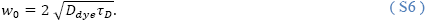

Here *D_dye_* is the diffusion coefficient for the fluorescent dye. The diffusion coefficient changes as a function of the temperature and needs to be corrected according to

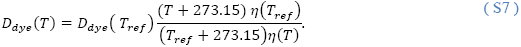

In Eq. (S6) all temperatures are in grad Celsius and *T_ref_* is a reference temperature for which the value of the diffusion coefficient is available. The *η(T)* is the dynamic viscosity at *T.* For water we have *η(27 °C)* = 0.8509 mPa sec and *η(37 °C)* = 0.6913 mPa sec (http://www.viscopedia.com/) yielding for the reference dyes Alexa488 and Alexa568 a mean diffusion coefficient of 463.23 μm^2^/sec and 521.46 μm^2^/sec, respectively.

## Supplementary Note 3: Correction for background and photobleaching

From fitting equation (S4) to the ACF of the fluorescent protein we extract the protein number *N*. The protein number needs to be corrected for photobleaching and background using the coefficients computed in *Fluctuation Analyzer*^1^

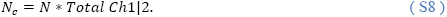

For low photon counts the computed bleach correction is less reliable. We found that for counts lower than twice the background counts in WT cells it is better to solely correct for the background

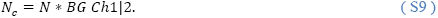

The correction parameters *Total Ch1* and *BG Ch1* as well as *Total Ch2* and *BG Ch2* for the FCS Channels 1 and 2, respectively are found in the result table from Fluctuation Analyzer. The concentration is calculated from the corrected number of proteins *N_c_* according to

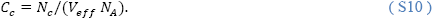

The effective volume *V_eff_* is computed from Eq. (S2) using the previously estimated values *κ* and *w_0_.* The computation of the corrected concentration is performed by *FCSFitM* and *FCSCalibration*.

## Supplementary Note 4: Estimate the number of proteins on small structures

The fluorescence intensity generated by a point-source of one fluorophore is defined by

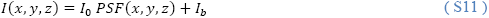

where *PSF(x,y,z)* is the confocal observation profile for the imaging settings, *I_0_* the fluorescence intensity characterizing a single fluorophore, and *I_b_* the background intensity. We denote the imaging volume by

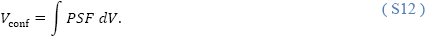

For FCS measurements performed within a volume larger than the PSF, we can assume a homogeneous distribution of fluorophores in space *I(x,y,z) =I*.For a linear detector the fluorescence intensity scales with the concentration *C* according to

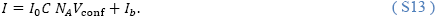

The concentration *C* is the concentration obtained from the FCS measurement and *N_A_* the Avogadro constant. We thus obtain the linear relationship

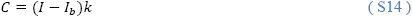

between concentration and fluorescence intensity with the calibration factor 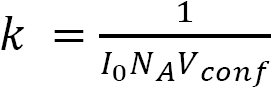.

An example calibration curve for simulated fluorophore distributions is shown in Supplementary Fig. 3 a-b. Relation (S14) holds true at every pixel with index *j*

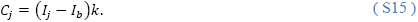

We can approximate the number of molecules at each pixel with

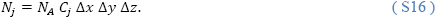

The parameters *Δx*, *Δy* and *Δz* characterize the pixel resolution in the 3 dimensions.

Equation (S16) can be applied at every pixel. To estimate the total number of proteins on a structure it is necessary to sum the protein number for all pixels in the structure of interest. For structures within the size of the PSF the signal must be integrated in 3D so that a large portion of the PSF is included. Simulations show that the estimation precision is directly proportional to the fraction of PSF included in the integration (Supplementary Fig. 3). Over 95% of the signal is accounted for when the region considered is more than twice the *e^2^* decay of the PSF (∼ 2/3 of the Airy disc diameter). If the size of the imaging PSF is known, approximations can be used to compute the signal from fewer pixels. For instance, a 3D Gaussian approximation of the PSF is obtained from the FCS measurement of the reference dye. These results can be used for the imaging PSF if the same imaging parameters are used (laser power and pinhole size). Below we provide some examples as a guideline.

## Supplementary Note 5: Total fluorophores in a point source

Consider *m* emitters concentrated at a point (Supplementary Fig. 3 c-d). The intensity is given by

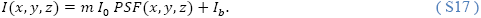

Using Eq. (S15) and after integration in 3D one obtains the expected value

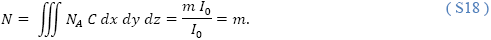

With (S15) and (S16) the integral is approximated by

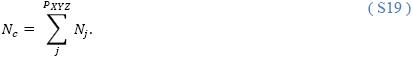

The sum is for all pixels that enclose the object of interest. When the parameters of the imaging PSF are known one can approximate Eq. (S18). Using the number of proteins in the plane of maximal intensity (here *z* = 0) one obtains

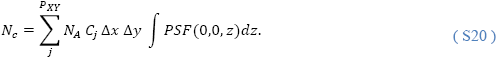

For a PSF approximated by a 3D Gaussian the Eq. (S20)

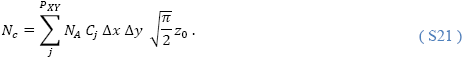

The peak intensity from the point source can also be used

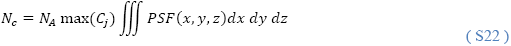

For a PSF approximated by a 3D Gaussian Eq. (S22) reads

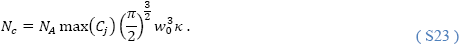

The smaller the imaging pixel size the more precise is the result (Supplementary Fig. 3 d, ≤ 0.6 *w*_0_).

## Supplementary Note 6: Density of fluorophores on a membrane

For a homogeneous spatial density the region size to be considered for quantification decreases due to symmetry properties of the membrane.

For example, for a membrane extending in the XZ-plane and parallel to the Y-axis we only need to consider pixels in one Z-plane (Supplementary Fig. 3 g). The density of fluorophores, *d_c_*, at a specific Y position is computed from the sum of pixels along the X direction in a region that enclose the fluorescence signal of the membrane (Supplementary Fig. 3 h, circles)

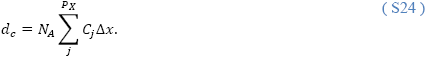

Using the peak intensity one obtains

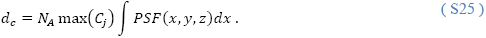

For a PSF approximated by a 3D Gaussian Eq. (S25) reads (Supplementary Fig. 3 h, squares)

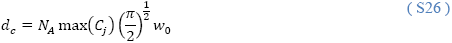

Similarly, the density *d_c_* of fluorophores on a membrane extending in the XY plane can be computed from the pixels in the Z direction (Supplementary Fig. 3 e). We denote 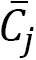 the average concentration in a specific Z-plane. One obtains

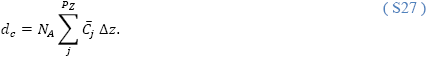

Using the plane peak intensity we have

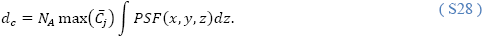

For a PSF approximated by a 3D Gaussian Eq. (S28) reads

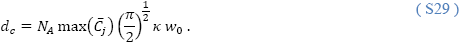

## REFERENCES

1. Cong, L., Ran, F., Cox, D., Lin, S. & Barretto, R. Multiplex Genome Engineering Using CRISPR / Cas Systems. Science 339, 819–823 (2013).

2. Shen, B. et al. Efficient genome modification by CRISPR-Cas9 nickase with minimal off-target effects. Nat Methods 11, 399–402 (2014).

3. Trevino, A. E. & Zhang, F. in Methods in Enzymology 546, 161–174 (Elsevier Inc., 2014).

4. Koch, B. et al. Generation and validation of homozygous fluorescent knock-in cells using genome editing. Nat. Protoc. (2018).

5. Digman, M. A., Stakic, M. & Gratton, E. in Methods in Enzymology 518, 121–144 (Elsevier Inc., 2013).

6. Krieger, J. W. et al. Imaging fluorescence (cross-) correlation spectroscopy in live cells and organisms. Nat. Protoc. 10, 1948–74 (2015).

7. Bacia, K., Kim, S. A. & Schwille, P. Fluorescence cross-correlation spectroscopy in living cells. Nat Methods 3, 83–89 (2006).

8. Bacia, K. & Schwille, P. Practical guidelines for dual-color fluorescence crosscorrelation spectroscopy. Nat. Protoc. 2, 2842–2856 (2007).

9. Elson, E. L. Fluorescence Correlation Spectroscopy: Past, Present, Future. Biophys. J. 101, 2855–2870 (2011).

10. Digman, M. A. & Gratton, E. Lessons in Fluctuation Correlation Spectroscopy. Annu. Rev. Phys. Chem. 62, 645–668 (2011).

11. Mahen, R. et al. Comparative assessment of fluorescent transgene methods for quantitative imaging in human cells. Mol. Biol. Cell 25, 3610–3618 (2014).

12. Walther, N. et al. A quantitative map of human Condensins provides new insights into mitotic chromosome architecture. bioRxiv 237834 (2018). doi:https://doi.org/10.1101/237834

13. Cuylen, S. et al. Ki-67 acts as a biological surfactant to disperse mitotic chromosomes. Nature 535, 308–312 (2016).

14. Germier, T. et al. Real-Time Imaging of a Single Gene Reveals Transcription-Initiated Local Confinement. Biophys. J. 113, 1383–1394 (2017).

15. Cai, Y. et al. An experimental and computational framework to build a dynamic protein atlas of human cell division. bioRxiv (2017). doi:doi.org/10.1101/227751

16. Conrad, C. et al. Micropilot: automation of fluorescence microscopy-based imaging for systems biology. Nat Methods 8, 246–249 (2011).

17. Tischer, C., Hilsenstein, V., Hanson, K. & Pepperkok, R. Adaptive fluorescence microscopy by online feedback image analysis. Methods Cell Biol 123, 489–503 (2014).

18. Wachsmuth, M. et al. High-throughput fluorescence correlation spectroscopy enables analysis of proteome dynamics in living cells. Nat Biotechnol 33, 384–389 (2015).

19. Germier, T. et al. Real-Time Imaging of a Single Gene Reveals Transcription-Initiated Local Confinement. Biophys. J. 113, (2017).

20. Kremers, G.-J., Gilbert, S. G., Cranfill, P. J., Davidson, M. W. & Piston, D. W. Fluorescent proteins at a glance. J. Cell Sci. 124, 157–160 (2011).

21. Shaner, N. C. et al. A bright monomeric green fluorescent protein derived from Branchiostoma lanceolatum. Nat. Methods 10, 407–409 (2013).

22. Shaner, N. C. et al. Improved monomeric red, orange and yellow fluorescent proteins derived from Discosoma sp. red fluorescent protein. Nat. Biotechnol. 22, 1567–1572 (2004).

23. Bindels, D. S. et al. mScarlet: A bright monomeric red fluorescent protein for cellular imaging. Nat. Methods 14, 53–56 (2016).

24. Ori, A. et al. Cell type-specific nuclear pores: a case in point for context-dependent stoichiometry of molecular machines. Mol Syst Biol 9, 648 (2013).

25. Suzuki, A., Badger, B. L. & Salmon, E. D. A quantitative description of Ndc80 complex linkage to human kinetochores. Nat. Commun. 6, 1–14 (2015).

26. Weir, J. R. et al. Insights from biochemical reconstitution into the architecture of human kinetochores. Nature 537, 249–253 (2016).

27. Bauer, M., Cubizolles, F., Schmidt, A. & Nigg, E. a. Quantitative analysis of human centrosome architecture by targeted proteomics and fluorescence imaging. Embo 35, 11–5 (2016).

28. Beck, M. et al. The quantitative proteome of a human cell line. Mol. Syst. Biol. 7, 549 (2011).

29. Zacharias, D. A., Violin, J. D., Newton, A. C. & Tsien, R. Y. Partitioning of Lipid-Modified Monomeric GFPs into Membrane Microdomains of Live Cells. Science 296, 913–916 (2002).

30. Cranfill, P. J. et al. Quantitative assessment of fluorescent proteins. Nat. Methods 13, 557–562 (2016).

31. Waters, J. C. Accuracy and precision in quantitative fluorescence microscopy. J. Cell Biol. 185, 1135 LP-1148 (2009).

32. Bancaud, A., Huet, S., Rabut, G. & Ellenberg, J. Fluorescence Perturbation Techniques to Study Mobility and Molecular Dynamics of Proteins in Live Cells: FRAP, Photoactivation, Photoconversion, and FLIP. Cold Spring Harb. Protoc. 2010, pdb.top90-top90 (2010).

33. Merzlyak, E. M. et al. Bright monomeric red fluorescent protein with an extended fluorescence lifetime. Nat. Methods 4, 555–557 (2007).

34. Lam, A. J. et al. Improving FRET dynamic range with bright green and red fluorescent proteins. Nat. Methods 9, 1005–1012 (2012).

35. Khmelinskii, A. et al. Tandem fluorescent protein timers for in vivo analysis of protein dynamics. Nat. Biotechnol. 30, 708–714 (2012).

36. Schwanhüusser, B. et al. Global quantification of mammalian gene expression control. Nature 473, 337–342 (2011).

37. Wu, J.-Q., McCormick, C. D. & Pollard, T. D. in Methods in cell biology 89, 253–273 (Elsevier Inc., 2008).

38. Verdaasdonk, J. S., Lawrimore, J. & Bloom, K. in Methods in Cell Biology 123, 347–365 (Elsevier Inc., 2014).

39. Wu, J.-Q. J.-Q. & Pollard, T. D. Counting Cytokinesis Proteins Globally and Locally in Fission Yeast. Science 310, 310–314 (2005).

40. Joglekar, A. P., Bouck, D. C., Molk, J. N., Bloom, K. S. & Salmon, E. D. Molecular architecture of a kinetochore-microtubule attachment site. Nat. Cell Biol. 8, 581–5 (2006).

41. Weidemann, T. et al. Counting Nucleosomes in Living Cells with a Combination of Fluorescence Correlation Spectroscopy and Confocal Imaging. J. Mol. Biol. 334, 229–240 (2003).

42. Maeder, C. I. et al. Spatial regulation of Fus3 MAP kinase activity through a reaction-diffusion mechanism in yeast pheromone signalling. Nat. Cell Biol. 9, 1319–1326 (2007).

43. Shivaraju, M. et al. Cell-cycle-coupled structural oscillation of centromeric nucleosomes in yeast. Cell 150, 304–316 (2012).

44. Ulbrich, M. H. & Isacoff, E. Y. Subunit counting in membrane-bound proteins. Nat. Methods 4, 319 (2007).

45. Ulbrich, M. H. in Springer Series on Fluorescence 13, 263–291 (2011).

46. Ta, H., Wolfrum, J. & Herten, D.-P. An extended scheme for counting fluorescent molecules by photon-antibunching. Laser Phys. 20, 119–124 (2010).

47. Ta, H. et al. Mapping molecules in scanning far-field fluorescence nanoscopy. Nat. Commun. 6, 7977 (2015).

48. Lawrimore, J., Bloom, K. S. & Salmon, E. D. Point centromeres contain more than a single centromere-specific Cse4 (CENP-A) nucleosome. J. Cell Biol. 195, 573–582 (2011).

49. Charpilienne, A. et al. Individual Rotavirus-like Particles Containing 120 Molecules of Fluorescent Protein Are Visible in Living Cells. J. Biol. Chem. 276, 29361–29367 (2001).

50. Picco, A., Mund, M., Ries, J., Nédélec, F. & Kaksonen, M. Visualizing the functional architecture of the endocytic machinery. Elife 4, (2015).

51. Keppler, A. et al. A general method for the covalent labeling of fusion proteins with small molecules in vivo. Nat. Biotechnol. 21, 86–89 (2003).

52. Gautier, A. et al. An Engineered Protein Tag for Multiprotein Labeling in Living Cells. Chem. Biol. 15, 128–136 (2008).

53. Los, G. V. et al. HaloTag: A Novel Protein Labeling Technology for Cell Imaging and Protein Analysis. ACS Chem. Biol. 3, 373–382 (2008).

54. Mütze, J., Ohrt, T. & Schwille, P. Fluorescence correlation spectroscopy in vivo. Laser Photonics Rev. 5, 52–67 (2011).

55. Lukinavicius, G. et al. SiR-Hoechst is a far-red DNA stain for live-cell nanoscopy. Nat Commun 6, 8497 (2015).

56. Neumann, B. et al. Phenotypic profiling of the human genome by time-lapse microscopy reveals cell division genes. Nature 464, 721–727 (2010).

57. Otsuka, S. et al. Nuclear pore assembly proceeds by an inside-out extrusion of the nuclear envelope. Elife 5, (2016).

58. Schindelin, J. et al. Fiji: an open-source platform for biological-image analysis. Nat Methods 9, 676–682 (2012).

59. Schindelin, J., Rueden, C. T., Hiner, M. C. & Eliceiri, K. W. The ImageJ ecosystem: An open platform for biomedical image analysis. Mol Reprod Dev 82, 518–529 (2015).

60. Rüttinger Buschmann, V., Krämer, B., Erdmann, R., MacDonald, R., & Koberling, F., S. et al. Comparison and accuracy of methods to determine the confocal volume for quantitative fluorescence correlation spectroscopy. J. Microsc. 232, 343–352 (2008).

61. Capoulade, J., Wachsmuth, M., Hufnagel, L. & Knop, M. Quantitative fluorescence imaging of protein diffusion and interaction in living cells. Nat. Biotechnol. 29, 835–839 (2011).

62. Kaputsa, P. Absolute diffusion coefficients: compilation of reference data for FCS calibration. (2010). Available at: https://www.picoquant.com/scientific/technical-and-application-notes/category/technical_notes_techniques_and_methods.

## References

1 Wachsmuth, M. et al. High-throughput fluorescence correlation spectroscopy enables analysis of proteome dynamics in living cells. Nature biotechnology 33, 384–389, doi:10.1038/nbt.3146 (2015).

2 Schwille, P. & Haustein, E. in Biophysics Textbook Online Vol. 1 1–62 (2001).

